# Neural dynamics outside task-coding dimensions drive decision trajectories through transient amplification

**DOI:** 10.1101/2025.11.20.689599

**Authors:** Ulises Pereira-Obilinovic, Kayvon Daie, Susu Chen, Karel Svoboda, Ran Darshan

## Abstract

Linking neural activity to behavior typically involves identifying activity subspaces that encode task information, such as stimuli, memory, and choices. However, it is unclear whether activity in these “coding dimensions” drives behavior or merely reflects the underlying computations. Neural activity in other “residual dimensions” is typically ignored. We developed a recurrent neural network that explicitly models interactions between coding and residual subspaces. Applied to multi-regional Neuropixels recordings from mice performing a delayed movement task, the model reveals that perturbations of residual dimensions reliably alter behavior, whereas perturbations of the choice dimension, which encodes the animal’s decision, are less effective. These effects arise because residual dimensions drive amplification across an intermediate number (∼10) of dimensions, before the dynamics settle into discrete attractors corresponding to choice. Our findings show that neural activity previously considered task-irrelevant can have critical roles in driving behavior.

## Introduction

During decision-making, neural activity reflects a wide range of behavioral variables. Much of the classical literature has centered on identifying neurons and neural populations with spike rates that correlate with task variables, such as sensory stimuli, memory, choices, or actions (Gold & Shadlen 2007), and such neurons are found across many brain areas (Steinmetz et al. 2019; Chen et al. 2024; Angelaki et al. 2025; Guo et al. 2014; Guo et al. 2017; Gao et al. 2018). Yet, correlations alone do not establish causal involvement. Although activating selective neurons can bias behavior (Salzman et al. 1990; Marshel et al. 2019; Robinson et al. 2020; Daie et al. 2021), recent work indicates that decisions can also be shaped by populations that exhibit little or no explicit tuning to the relevant task variables (Daie et al. 2023; Gauld et al. 2024; Bounds et al. 2024).

The advent of methods for large-scale neural recordings (Ahrens et al. 2013; Sofroniew et al. 2016; Jun et al. 2017) has shifted the focus from individual neurons to the geometry of population activity (Churchland et al. 2007; Perich et al. 2025). Dimensionality-reduction techniques (Cunningham & Yu 2014; Pang et al. 2016; Kobak et al. 2016; Williams et al. 2018; Li et al. 2016; Pellegrino et al. 2024) reveal low-dimensional “coding” subspaces that capture most task-related variance. The remaining “residual” dimensions exhibit weak or heterogeneous task modulation. These methods have provided influential descriptions of neural trajectories during decision-making (Vyas et al. 2020; Gallego et al. 2017; Li et al. 2016; Steinemann et al. 2024; Monsalve-Mercado & Miller 2025; Mante et al. 2013; Inagaki et al. 2019). The prevailing interpretation is that behaviorally relevant dynamics unfold within the coding subspace, including maintenance of short-term memories, decision-making, motor plans and actions. Activity in residual dimensions is treated as noise or an epiphenomenon. However, just as tuning in single neurons does not guarantee causal relevance, it remains unclear whether the extracted coding dimensions drive decisions—or merely correlate with them.

Residual dimensions show weak encoding of task variables. Can they play roles in shaping task-related population dynamics? We developed a neural network framework that explicitly separates task-coding and residual subspaces. The model includes all recurrent interactions between these subspaces, while constraining the coding dimensions to represent key task variables (sample, choice, response) and defining residual dimensions as orthogonal directions capturing the remaining variance. Unlike standard dimensionality-reduction approaches, this framework allows us to model how interactions between coding and residual dimensions influence neural trajectories at the single-trial level. Compared to previous activity-constrained network models (Rajan et al. 2016; Finkelstein et al. 2021; DePasquale et al. 2023; Valente et al. 2022; Kim et al. 2023; Sourmpis et al. 2023; Pals et al. 2024; Sourmpis et al. 2024; Langdon & Engel 2025), our approach focused on the low-dimensional latent variables governing both sets of dimensions, identify how they are dynamically coupled, while decoupling their distinct contribution to neural computation.

We applied this framework to large-scale recordings from mouse anterior lateral motor cortex (ALM) and connected thalamus during a delayed movement task (Chen et al. 2024). Model predictions showed that perturbations directed along *residual* dimensions—orthogonal to the choice axis— disrupt neural trajectories and bias decisions, whereas perturbations along *coding* dimensions are much less potent (Daie et al. 2023). Neural dynamics unfolded within subspaces of intermediate dimensionality (∼10 dimensions), substantially higher than the low-dimensional coding subspace. Within these subspaces, trajectories evolved through non-normal amplification (Goldman 2009; Murphy & Miller 2009): interactions between nearly orthogonal activity patterns transiently amplified small deviations before converging onto discrete attractor states corresponding to behavioral choice (Inagaki et al. 2019; Oña-Jodar et al. 2024). As a result, components in the residual subspace that are weakly tuned and explain little activity variance can grow transiently and redirect trajectories within the coding subspace. Our results highlight the importance of population dynamics outside task-aligned dimensions.

## Results

### Coding and residual dimensions in activity-trained networks

To investigate how residual dimensions contribute to neural computations, we developed a recurrent neural network (RNN) model trained to simultaneously fit large-scale neural recordings and uncover latent subspaces associated with behavior. We distinguished between coding dimensions—subspaces aligned with task-relevant variables such as sensory stimuli, choices, and motor outputs—and residual dimensions, which are orthogonal to the coding subspace but may still contribute to overall neural dynamics (Fig. 1a).

**Figure 1:**
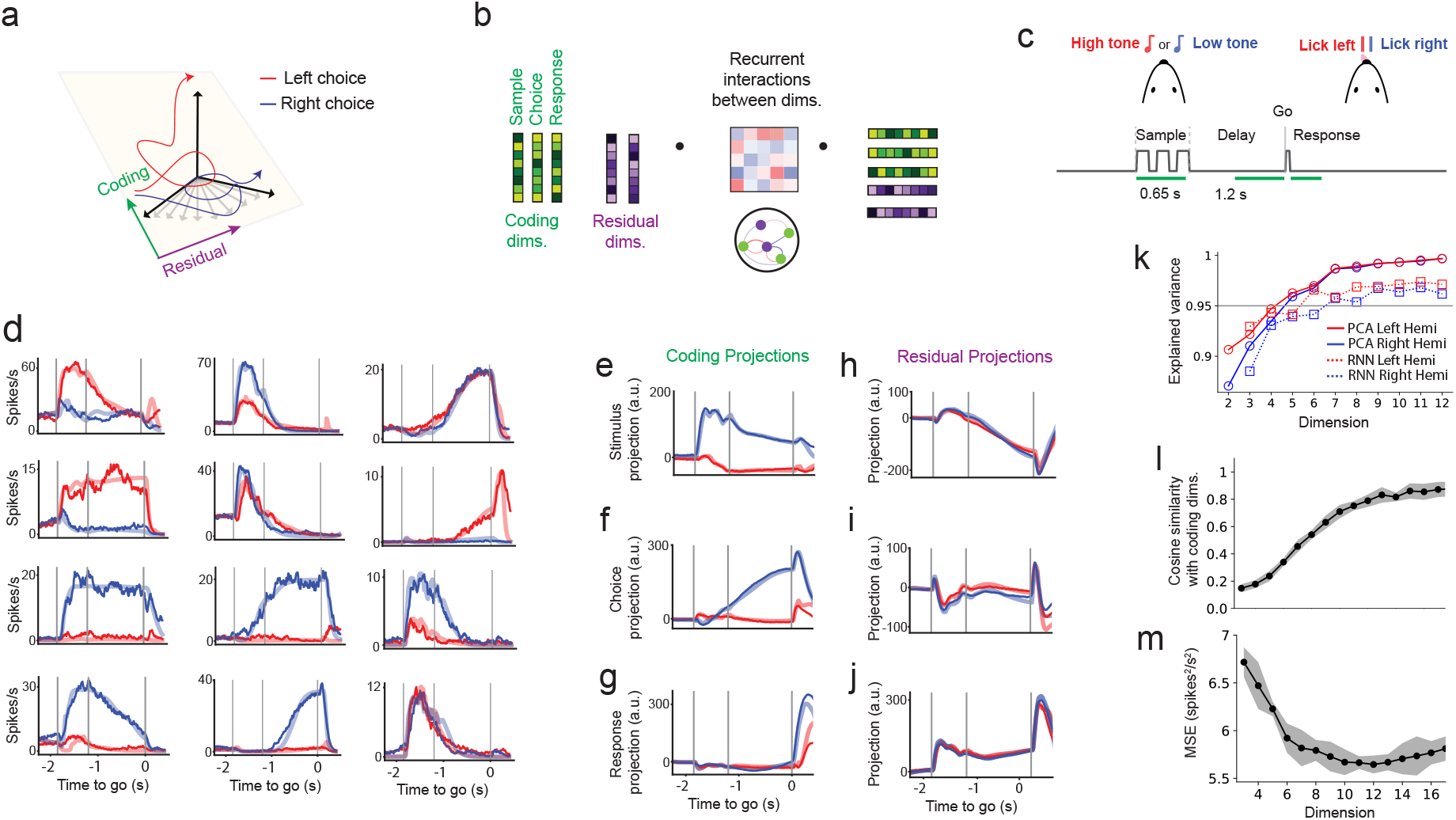
Coding and residual dimensions in activity-trained networks, applied to a memory-guided decision-making task. (a) Neural trajectories during decision-making corresponding to left and right choices. (b) Network connectivity is defined by all possible combinations of coding and residual dimensions. The vectors representing these dimensions and their recurrent interactions are optimized jointly to fit neural activity. Coding dimensions are constrained to encode behavioral variables. (c) Memory-guided movement task. Auditory stimuli (high tone or low tone) presented during the sample epoch instruct a directional movement (lick left or right), after a 1.2 s delay epoch. Green bars indicate the windows for calculating coding dimensions: first 600 ms of auditory stimulus during the sample epoch; last 600 ms of the delay epoch; first 350 ms post Go cue. (d) Trial-averaged activity for correct left and right trials (ALM, right hemisphere). Dark lines, neural recordings; light lines, model predictions. Total number of dimension, *P* = 10. (e-g) Projections of neural activity onto coding dimensions. Projections align with task variables stimulus, choice, and response during the sample, delay, and response epochs (c). (h-j) Projections, residual dimensions. Projections contain little selective activity. (k) Explained variance of neural activity from PCA (circles, solid lines) and RNN (squares, dashed lines) as a function of the number of dimensions. (l) Average cosine similarity between the coding dimensions extracted from the network fit and those derived from the data shown as a function of model dimensionality. Shaded regions indicate the standard error computed across five independent training sets using 80% of the trials. (m) Cross-validated mean squared error (MSE) between the network’s firing rates and the trial-averaged activity computed on the held-out 20% of trials in each subset. Error bars indicate the standard error across folds (see Methods). In (l) and (m), cosine similarity and MSE curves are averaged across networks trained separately on each hemisphere; hemisphere-specific results are shown in Fig. S1.

We modeled the firing-rate dynamics of each recorded neuron, *r*_*i*_(*t*), using a nonlinear recurrent neural network of *N* units, whose activity evolves under recurrent interactions specified by the connectivity matrix **J** (Methods, Eq. (3)). Each neuron received external input, representing task events such as the sensory stimulus during the sample epoch and a go cue (Fig. 1c). Neuronal dynamics were governed by the standard rate equation (Grossberg 1969; Hopfield 1984; Miller & Fumarola 2012). Each neuron transformed its net input through a nonlinear transfer function, ϕ_*i*_(*x*), with heterogeneous saturation levels across the population (see Methods, Eq. (4)).

In addition, the RNN explicitly captured interactions between coding and residual dimensions. The connectivity matrix between neurons (*i, j* ∈ {1…*N*}) was constrained to be low-rank (Mastrogiuseppe & Ostojic 2018):

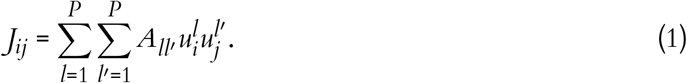

The low-rank structure was further defined by a set of orthonormal vectors: 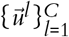 spanning the *C* coding dimensions (Methods and see below) and 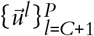 spanning the residual subspace, which is orthogonal to the coding subspace.

The interaction matrix ***A*** governs how population dynamics within and across these subspaces interact (Fig. 1b). This formulation can capture a broad range of population dynamics, including fixed-point attractors (Hopfield 1982), sequential activity (Kleinfeld 1986; Sompolinsky & Kanter 1986; Gillett et al. 2020; Gillett & Brunel 2024), and continuous attractors (Ben-Yishai et al. 1995; Seung 1996; Darshan & Rivkind 2022; Khona & Fiete 2022). By explicitly parameterizing interactions between coding and residual dimensions in the interaction matrix ***A***, we are able to discover and dissect the interactions between task-coding and residual components of neural dynamics.

Projections of the neural trajectories,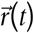, onto the coding and residual subspaces define the latent variables in our model

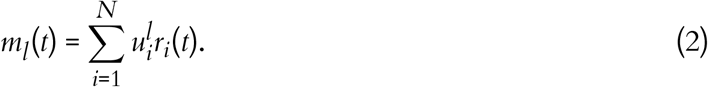

By projecting the network dynamics onto the coding and residual subspaces, we obtained a reduced description of the population activity in terms of latent variables 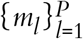, which correspond to the projection of firing rates onto the low-rank basis (see Methods, Eq. (11)). The latent dynamics evolve according to a *P*-dimensional nonlinear dynamical system that captures how population activity evolves within the coding and residual subspaces. This formulation enables us to assess the extent to which residual dynamics shape trajectories along coding dimensions.

### Training on neural recordings from the anterior lateral motor cortex (ALM)

We applied our modeling framework to large-scale electrophysiological recordings spanning cortical and subcortical structures in mice performing a memory-guided movement task (Chen et al. 2024) (Fig. 1c; see Methods). On each trial, a brief auditory cue (either high or low frequency) instructed a left or right lick response following a 1.2-second delay. This task requires the animal to transform a transient sensory input into a persistent motor plan maintained over the delay epoch (Li et al. 2016). We refer to trials in which the animal’s lick direction matched the instructed stimulus as correct trials; unless stated otherwise, all analyses use correct trials only. Error trials, in which the animal licked in the direction opposite to the instructed stimulus, are introduced in the single-trial section

To identify the *C*-dimensional subspace aligned with task-relevant variables, we used neural activity averaged across trials. In the resulting population trajectories, each time point is a point in an *N*-dimensional activity space (Georgopoulos et al. 1988; Stopfer et al. 2003). We used dimensionality reduction (Methods) to identify the directions that best separated behavioral conditions, such as left versus right choices. This approach isolates interpretable task-relevant dimensions of population activity (Li et al. 2016; Inagaki et al. 2018). We focused on three dimensions: sample, choice, and response (*C* = 3). The sample dimension captured stimulus-evoked activity in the first 600 ms of the trial; the choice dimension separated left and right trials during the delay epoch (600 ms before the go cue); and the response dimension distinguished left and right licks 350 ms after the go cue (Li et al. 2016; Chen et al. 2024) (Fig. 1c). The sample, choice, and response coding dimensions explain approximately 47%, 58%, and 43% of the trial-average variance during the sample, delay, and response epochs, respectively (see Methods), consistent with previous reports (Inagaki et al. 2018). These contain variance related to the core computational demands of the task: sensory integration (Finkelstein et al. 2021), choice maintenance (Li et al. 2016; Chen et al. 2021; Wang et al. 2021), and motor execution (Inagaki et al. 2022; Chen et al. 2024). Among these dimensions, the choice dimension has received particular attention (Li et al. 2016; Finkelstein et al. 2021; Inagaki et al. 2022; Daie et al. 2023; Wang et al. 2021; Chen et al. 2024), as it is strongly correlated with the animal’s upcoming action (Inagaki et al. 2019) and remains stable throughout the delay epoch (Li et al. 2016; Inagaki et al. 2018; Daie et al. 2023).

The remaining *R* = *P*–*C* dimensions define the residual subspace. The residual dimensions are orthogonal to the coding dimensions. We allowed the RNN to discover how they support the observed dynamics throughout the training procedure. The training of the RNN thus jointly identifies the low-dimensional subspace spanned by both coding and residual dimensions, the interaction structure among these dimensions, and the recurrent dynamics that best reproduce the neural data.

We first focused on ALM, which is required for maintaining choice-related activity (Guo et al. 2014) and movement initiation (Inagaki et al. 2022). We trained the network to reproduce condition-averaged activity from 7,884 neurons across mice(3,640 in the left hemisphere, 4,244 in the right) during correct left and right trials (Fig. 1d). During the sample epoch (Fig. 1c), the network received two distinct inputs corresponding to left and right auditory cues that were projected onto both task-coding and residual dimensions (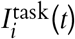 in Eq. (7,8), Methods). The alignment of these inputs was optimized through training (see Methods). A similar input was received by the network for the go cue (Eq. (7), Methods).

The network integrates transient stimuli during the sample epoch to generate and maintain preparatory activity during the delay epoch, and the go cue subsequently reshapes this preparatory activity to produce movement-related dynamics (Eq. (7), Methods). By explicitly separating coding and residual dimensions with associated latent variables, the model provides a framework for examining how sensory inputs are integrated by the recurrent dynamics along specific latent dimensions to generate neural dynamics that match ALM population activity. Projections onto task-coding dimensions recapitulated known features of ALM activity (Fig. 1e–g), including ramping along the choice axis during the delay epoch (Li et al. 2016; Chen et al. 2024) (Fig. 1f). Residual dimensions exhibited little selectivity for left versus right conditions (Fig. 1h–j).

Trial-averaged neural activity resides in a low-dimensional subspace (Inagaki et al. 2018). Principal component analysis (PCA) of the trial-averaged population responses revealed that over 95% of the variance was captured by the first five components (Fig. 1k). Our trained low-rank models reflected a similar low-dimensional structure. To quantify this, we first fixed the model’s three components to align with empirically estimated coding dimensions and trained only the recurrent interaction matrix. This configuration explained 78% of the variance. Next, we introduced a soft alignment constraint, allowing the coding components and interaction matrix to be co-optimized, while penalizing deviations from the data-derived coding dimensions (Methods). This increased the explained variance to nearly 90%.

Despite explaining a large proportion of the variance, models restricted to three dimensions with the soft alignment constraint exhibited poor alignment with the empirical task-coding axes (Fig. 1l). This indicated that three dimensions were insufficient to simultaneously reproduce neural activity and align with the task-coding subspace. We therefore retained the soft alignment and incrementally incorporated additional residual dimensions. Adding these dimensions yielded only a modest increase in explained variance (from 90% to 97%; Fig. 1k), but this small variance gain was accompanied by a substantial improvement in alignment with the empirical task-coding axes (Fig. 1l and Fig. S1a). The improved alignment plateaued around 12 total dimensions. Thus, a combination of coding and residual dimensions are needed in the network model to capture both the task-relevant and single neuron dynamics.

Notably, residual dimensions individually account for only 0.1–0.2% of the trial-average variance (see Methods), compared to 47%, 58%, and 43% explained by the coding dimensions during the sample, delay, and response epochs, respectively.

We focus subsequent analyses on a 10-dimensional model (Figs. 2–4), which provided a good fit (Fig. 1m) while maximizing alignment with data-derived coding axes (Fig. 1l, m; Fig. S1a,b). All the results below were robust across intermediate dimensionalities (*P* = 9–13), multiple random seeds, and both recorded hemispheres.

**Figure 2:**
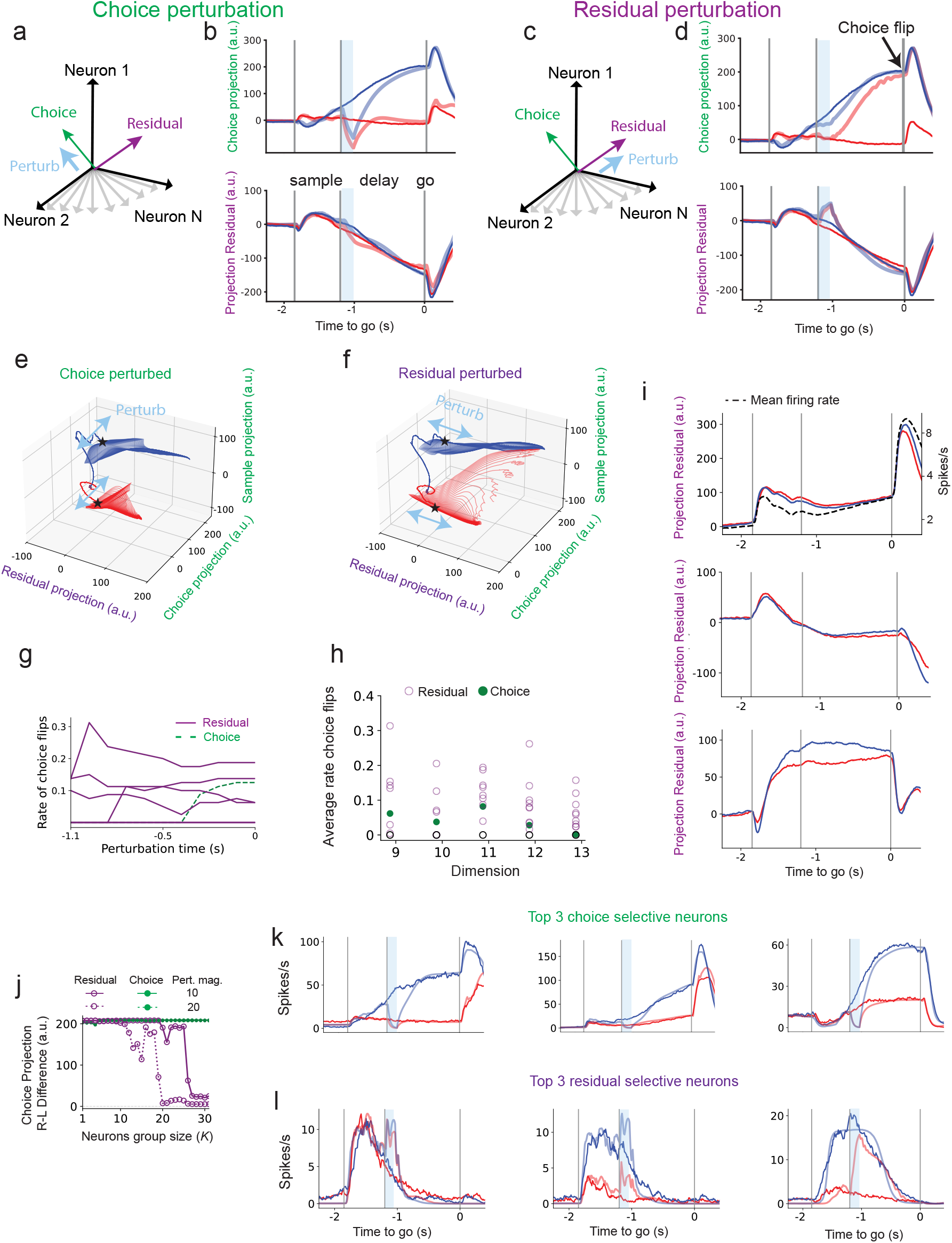
Choice-aligned perturbations preserve choice, whereas residual-aligned perturbations disrupt it. (a) Network perturbation in the choice direction (blue arrow). (b) Projections of neural activity (Dark hues) and network dynamics (light hues) onto the choice (top) and residual (bottom) dimensions for perturbations aligned with the choice dimension. Blue bar, the duration of the network perturbation. (c-d) Same as (a-b), but for perturbations aligned with one of the residual dimensions that led to a different endpoint of the choice-projected trajectory (‘influential residual’). (e-f) Network responses for perturbations with magnitudes ranging from –10 to 10 applied along the choice dimension (e) and an influential residual dimension (f) indicated by blue arrows, visualized in the 3D subspace defined by choice, sample, and residual axes. Trajectories are shown until the go cue. Black stars, start of the delay epoch.(g) Rate of choice flips (the fraction of perturbations crossing a predefined choice projection threshold; see panels e and f) for 11 distinct 100 ms perturbation intervals applied during the delay epoch, across 10 network dimensions. Each dimension was tested with perturbation magnitudes from –15 to 15 (see Methods). (h) Delay epoch average choice-flip rates from (g), summarized across networks trained with 9 to 13 dimensions. Black circles indicate the average effect of 10 random perturbations orthogonal to both the coding and residual dimensions, matched in perturbation magnitude. (i) Dynamics of projections onto three additional influential dimensions beyond the one shown in (d). Neural activity is projected onto each influential dimension. In the top panel, the dashed line denotes the mean firing rate averaged across neurons and left/right trials. (j) Magnitude of the difference in the choice projection between left and right trials, averaged over the 100 ms preceding the go cue, as a function of perturbed group size. For each group size *K* (number of neurons in the perturbed set), we perturb the top-*K* neurons ranked by selectivity, either the top-*K* choice-selective or the top-*K* residual-selective neurons (Methods). Perturbations are applied during the first 100 ms of the delay epoch as in (a-d). (k) Responses of the top 3 choice-selective neurons to perturbations targeting the top 30 choice-selective neurons. (l) Same as (k), but for the top 3 residual-selective neurons when targeting the top 30 residual-selective neurons.

Having established a network model that accurately captures both single-neuron activity and latent dynamics along the coding and residual subspaces, we next asked: what computational role do these residual dimensions play in shaping decision-related dynamics in this task?

### Choice-aligned perturbations preserve choice, whereas residual-aligned perturbations disrupt it

We next probed the trained model with targeted perturbations of its activity. As the projection on choice at the go cue predicts the animal’s movement direction, we consider a flip of the trajectory at the end of the delay epoch as a flipped decision (Li et al. 2016; Inagaki et al. 2019). Perturbations along the choice dimension (Fig. 2a), despite its strong correlation with behavior, had little effect on choice (Fig. 2b; Fig. S2 a-c). Indeed, projecting the network dynamics onto the sample, choice, and residual dimensions revealed that perturbations along the choice dimension caused only transient deviations, which rapidly decayed back to the unperturbed trajectory (Fig. 2e). In contrast, perturbing a subset of residual dimensions, which have minimal choice selectivity, triggered large and sustained excursions and consistently flipped the choice projection from one choice to the other (e.g., left to right in Fig. 2d, Fig. 2f; Fig. S2 d-i). These effects were robust across hemispheres (Fig. S2j-n).

Analysis of the network’s response to perturbations (see Methods) revealed that a subset of residual dimensions, when perturbed during the delay epoch, consistently showed a pronounced likelihood of inducing choice flips (switches of the endpoints of choice-projected trajectories;Fig. 2g; Fig. S2n). We termed these dimensions *influential residual* dimensions. In contrast, perturbations to the choice dimension are largely ineffective early in the delay epoch (Fig. 2g). Although perturbations to the choice dimensions become more effective later, they never catch up to the effectiveness of residual dimensions. This temporal progression is consistently observed across multiple trained networks and different numbers of residual dimensions, *R* = 10 – 13 (Fig.S2o). Perturbations constructed to lie in the subspace orthogonal to the coding and residual dimensions had no effect on choice (Fig. 2h and Fig. S2o, black markers), indicating that perturbations affect choice only when acting within the coding–residual subspace. Overall, our analyses show that residual dimensions are generally more influential in perturbing choice than the choice dimension (Fig. 2g and Fig. S2n).

Some of the influential residual dimensions correlate with population activity patterns previously associated with decision-making and motor preparation, such as ramping activity (Thura & Cisek 2014; Tanji & Evarts 1976; Churchland et al. 2010; Guo et al. 2014),(Fig. 2d, bottom) or the trial averaged activity across trials (Fig. 2i, top). Other dimensions show no correspondence with task-related signals (Fig. 2i, middle and bottom). Moreover, residual dimensions exhibit little or no choice selectivity compared to the coding dimensions (Fig. 1l). But some of these influential residual dimensions show non-selective ramping (e.g., Fig. 1h) which have been previously associated with the maintenance of prospective motor plans during the delay epoch (Li et al. 2016; Inagaki et al. 2019; Finkelstein et al. 2021). These observations suggest that a dimension’s influence on choice is not necessarily tied to its alignment with task-evoked activity or choice selectivity.

Experimental perturbations act on subsets of neurons (Marshel et al. 2019; Daie et al. 2021; Adesnik & Abdeladim 2021; Daie et al. 2023), and their behavioral effects are determined by how these neurons project onto the coding and residual subspaces. A natural strategy for influencing behavior is therefore to perturb the most choice-selective neurons. However, our results show that this intuition can fail when influential dimensions lie within the residual subspace. To test this, we perturbed the top-*K* neurons most aligned with either the choice dimension or an influential residual dimension to test whether few-cell perturbations disrupt choice. Choice-selective neurons have higher mean firing rates during the delay epoch (14 vs. 4 spikes/s; see Fig. S2p vs. q, continuous traces) and substantially stronger delay-epoch choice tuning (15 vs. 0.3 spikes/s, right–minus–left; dashed traces in Fig. S2p vs. q). Despite these larger firing rates and stronger choice selectivity, perturbing the top-*K* choice-aligned neurons had little effect on behavior. In contrast, perturbing the top-*K* residual-aligned neurons reliably flipped the network’s choice. Indeed, perturbing approximately 20–30 of the most residual-selective neurons abolished the left–right separation on the choice dimension, flipping the network’s choice (Fig. 2j-l). By contrast, perturbing the top choice-selective neurons transiently reduced choice selectivity during the 100-ms perturbation, but coding rebounded once the input ceased (Fig. 2j–k; See also Fig. S2r–t for opposite-sign excitatory perturbations).

### Non-normal interactions between coding and residual dimensions

The interaction matrix, **A**, organizes the flow of activity across subspaces. (Eq. (11)). Therefore, to uncover the mechanisms by which residual dimensions exert control over choice dimension we analyzed the structure of **A**.

We discovered strong, asymmetric interactions between coding and residual dimensions (Fig. 3a), implying that decision-making arises not solely from task-aligned neural signals– as is often assumed in models of decision-making and short-term memory (Wang 2002; Inagaki et al. 2019). Interactions between coding dimensions and residual dimensions are critical.

**Figure 3:**
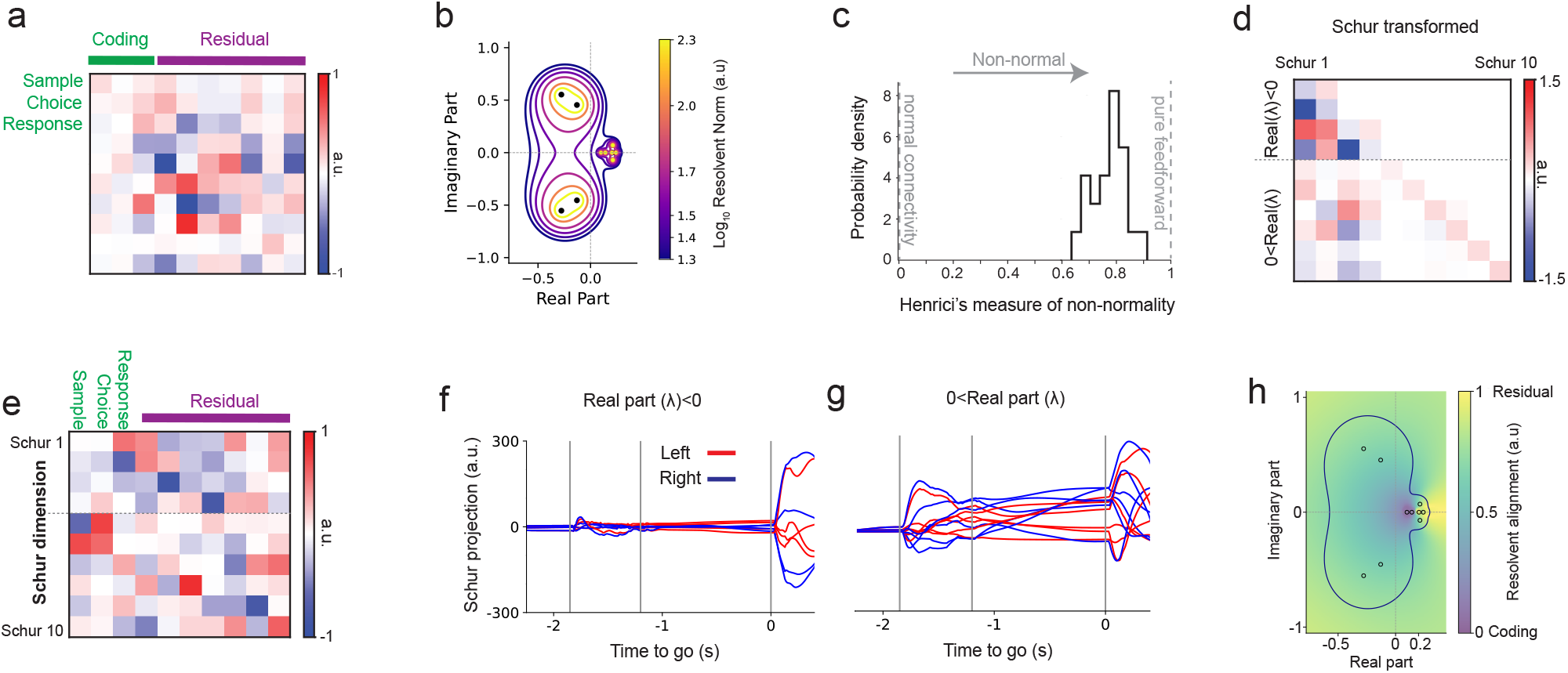
Non-normal interactions between coding and residual dimensions shape decision trajectories. (a) Interaction matrix **A** for the network shown underlying the data in Fig. 2 (latent dimensionality *P* = 10. (b) Eigenvalue spectrum and pseudospectra of the interaction matrix (Trefethen & Embree 2020). Contour lines correspond to resolvent norms of 5 × 10^−2^, 4 × 10^−2^, 3 × 10^−2^, 2 × 10^−2^, 1 × 10^−2^, and 5 × 10^−3^ (Methods). (c) Normalized Henrici index (see Methods) across 40 trained networks with latent dimensionality ranging from *P* = 9 to *P* = 16. (d) Schur form (lower-triangular matrix) of **A**. Diagonal entries indicate the real parts of eigenvalues; red and blue indicate Schur dimensions with positive and negative real parts, respectively. Dashed line separates the two groups. (e) Unitary matrix defining the orthonormal Schur basis. Rows correspond to Schur vectors; the dashed line separates dimensions associated with positive versus negative real eigenvalues. (f, g) Network activity projected onto Schur dimensions associated with negative (f) and positive (g) real eigenvalues for left (red) and right (blue) choice trials. (h) Alignment of the resolvent to the coding and residual subspaces. Alignment between the dominant right-singular vector of the resolvent and the residual subspace (see Methods). Yellow indicate regions in the complex plane where the most amplifiable input directions align strongly with residual subspace, whereas purple regions indicate strong alignment with the coding subspace. The lowest resolvent contour from panel b is overlaid for visual correspondence of both panels.

In the trained network, perturbations to some of the residual dimensions at the beginning of the delay epoch were amplified and influenced the rate of choice flips later during the delay epoch. We hypothesized that this amplification reflects a network mechanism in which non-normal interactions (Ganguli et al. 2008; Goldman 2009; Murphy & Miller 2009; Hennequin et al. 2012) couple the coding and residual subspaces, allowing perturbations confined to residual directions to grow and ultimately impact choice. A non-normal connectivity matrix describes a network where activity can be temporarily amplified or suppressed due to asymmetries in how the different dimensions influence each other, even if the overall activity eventually settles down. This means that certain patterns of input can trigger strong, transient responses that would not occur in a network with symmetric (normal) connectivity.

We assessed the degree of non-normality of the interactions using pseudospectra analysis (Trefethen & Embree 2020), a method that expands upon eigenvalue analysis by quantifying the network’s sensitivity to perturbations. Standard eigenvalue analysis identifies directions in neural state space along which activity grows (positive eigenvalues) or fades (negative eigenvalues). However, this approach can miss important effects in networks with non-normal interactions, where the eigenvalues can be highly sensitive to small variations in the interactions. Pseudospectral analysis (see Methods) reveals this hidden sensitivity and identifies non-normal interactions. For networks with normal interactions, pseudospectra analysis shows isolated contours encircling eigenvalues. In our models, the contours were broad and overlapped multiple eigenvalues, especially around areas linked to amplifying activity (positive eigenvalues), which is a clear sign of strong asymmetric and non-normal interactions (Fig. 3b; Fig. S3a-e).The trained networks consistently exhibited strong non-normality (mean normalized Henrici index (Trefethen & Embree 2020; Asllani et al. 2018) across networks: 0.78; 0 for normal matrices and 1 for strictly non-normal; Fig. 3c; Fig. S3f).

We used a Schur decomposition (Methods) to divide the interaction matrix into two components: normal (symmetric) recurrent interactions and non-normal (asymmetric) interactions (Golub & Van Loan 2013). The Schur decomposition provides insight into how transient neural activity emerges from non-normal amplification (Goldman 2009; Murphy & Miller 2009; Hennequin et al. 2012) (Fig. 3d,e). We identified two distinct sets of dimensions: decaying Schur dimensions with negative eigenvalues and transiently amplifying Schur dimensions with positive eigenvalues. Decaying Schur dimensions exhibited large projection magnitudes during the response epoch (Fig. 3f), whereas amplifying Schur dimensions exhibited large projection magnitudes predominantly during the sample epoch and early in the delay epoch, and persisted during the delay epoch (Fig. 3g). Perturbations to amplifying Schur dimensions (positive eigenvalues) strongly impacted decision outcomes early in the delay epoch (Fig. S3g, red curves), underscoring their causal role in shaping decision-related neural dynamics. Conversely, perturbations of decaying Schur dimensions (negative eigenvalues) (Fig. S3g, blue curves) had pronounced effects mainly towards the go cue (Fig. S3h-j). Together, these results support distinct, epoch-specific roles for Schur dimensions, with transiently amplifying Schur dimensions contributing causally to delay-epoch activity and decision outcomes, and decaying Schur dimensions contributing primarily during the response epoch. Since the Schur decomposition expresses each dimension as a linear combination of coding and residual dimensions (Fig. 3e), this analysis shows that residual dimensions—despite little correlation with task variables—play a critical role in shaping task-related neural dynamics.

To assess the role of residual dimensions in non-normal amplification, we identified the latent dimensions that the network transiently amplifies and examined how they align with coding and residual subspaces in the fitted networks (Methods). Regions of strongest amplification, located near eigenvalues with positive real parts and reflecting shared non-normal interactions among latent dimensions (Fig. 3b), consistently aligned with residual dimensions (Fig. 3h, compare with Fig. 3b; see Methods). These eigenvalues correspond to Schur dimensions that exert strong effects on flipping the choice during the delay epoch (Fig. S3g). These results show that the strongest non-normal transient amplification in latent space, induced by the interaction matrix **A**, predominantly unfolds along residual directions.

A key question is whether non-normal connectivity is necessary to account for the data, or whether it represents just one of many solutions that fit the neural activity equally well. To test this, we constrained the network to have symmetric connectivity (Methods; Fig. S4k), it achieved comparable reconstruction of trial-averaged activity but produced qualitatively opposite perturbation responses: choice perturbations were strongly effective while residual perturbations were largely ineffective (Fig. S4l), the reverse of what we observed experimentally (Daie et al. 2023). Analysis of the network dynamics revealed that the choice dimension aligned with eigenvectors of near-zero eigenvalues of the interaction matrix (Fig. S4m), indicating that the network learned an approximate integrator along the choice dimension (Fig. S4n). These results demonstrate that reconstruction quality alone is insufficient to distinguish between mechanistically distinct circuits, and that non-normal interaction structure is necessary to simultaneously account for the neural activity and the causal perturbation responses.

Our results illustrate a critical distinction between dimensions that are optimal for decoding task variables and those dynamically influential in controlling neural trajectories. Although the choice dimension effectively encodes decision outcomes, influential residual dimensions are aligned with the transiently amplifying subspace, emphasizing the role of dynamics in the residual subspace in the transient non-normal dynamics that shapes decision trajectories.

### Dynamics during the delay epoch terminates in attractors corresponding to left and right choices

Previous analysis of the responses of ALM neuron populations to transient perturbations during the delay epoch has provided support for two disparate mechanisms: discrete fixed points related to choice (Inagaki et al. 2019; Finkelstein et al. 2021) and transient feed-forward dynamics (Daie et al. 2023).

Our model reconciles these two mechanisms. We examined the dynamics of the trained network at the end of the delay epoch. The network was trained on neural dynamics with a fixed delay epoch (1.2 s), terminated by a go cue. We withheld the go cue from the model after training. The network settled into one of two stable attractor states, each representing a distinct choice (Fig. 4a). Similarly, in the recordings, in a small subset of trials, the go cue was delayed from 1.2s to 1.8s (Methods). Importantly, these trials were not used for training the networks. Simulating these catch trials in our network by delaying the go cue by 0.6s revealed that by 1.2s, the network reached a choice-selective attractor. The late go cue at 1.8s collapses the selective attractor states back to a non-selective state (Fig. 4b). Consistent with the network’s dynamics, a neuron-subsampling bootstrap analysis of the catch trials (Methods) showed that for right trials during the original delay period the choice-projected activity ramps with a positive slope (mean slope = 0.23, *p* < 10^−3^). However, during the catch period the activity stabilizes with a slope not significantly different from zero (mean slope = –0.01, *p* = 0.86) (Fig. 4c; green), with the decrease in slope being significant across bootstrap iterations (*p* < 10^−3^; orange) (Maimon & Assad 2006; Tanaka 2007; Inagaki et al. 2019). To assess the dynamical stability of the fixed points, we computed the eigenvalues of the Jacobian of the dynamics after the go cue (see Methods) and found that the eigenvalue with the largest real part was negative across trained networks (Fig. 4d), confirming that the dynamics settled into one of the two choice-selective stable attractor states. Our results were also replicated in networks trained on recordings from the left hemisphere (Fig. S4).

**Figure 4:**
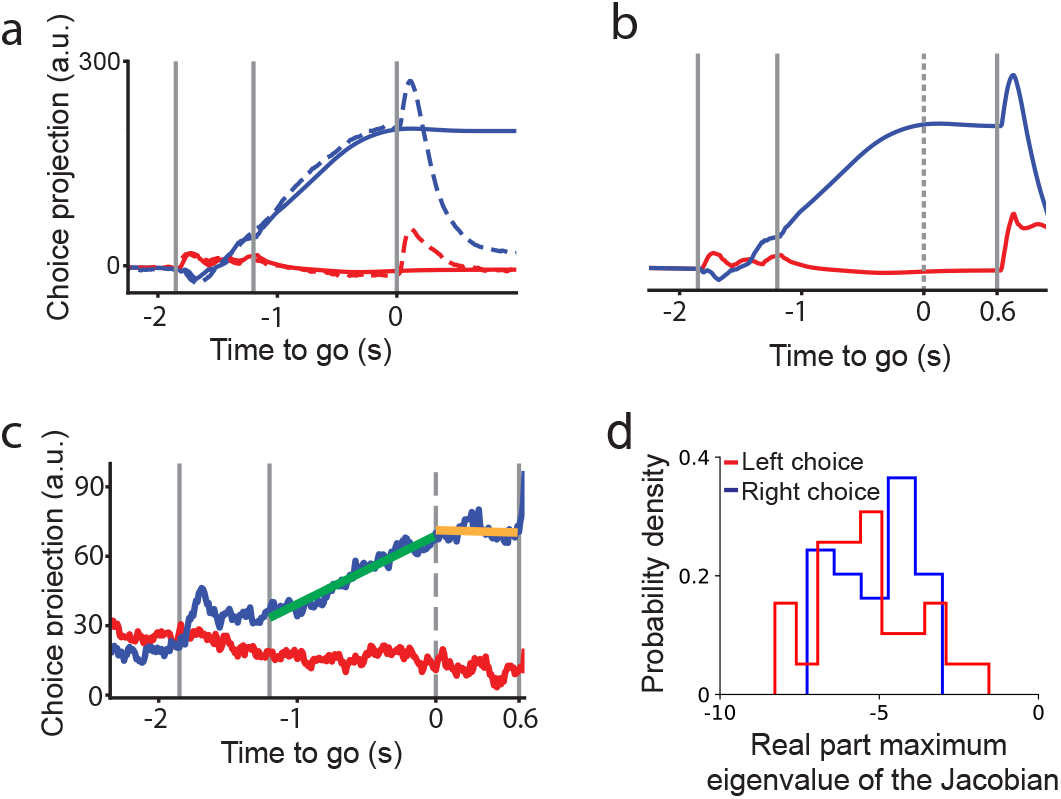
Dynamics during the delay epoch terminates in attractors corresponding to left and right choices. (a) Choice projections. Dashed lines, neural recordings for 1.2s delay; solid lines,network when the go cue is withheld. Choice projection from the network when the go cue is presented after a 1.8s delay epoch, simulating catch trials. The standard go cue used for training is indicated by a dashed line, corresponding to a 1.2 s delay. (c) Trial-averaged activity (2529 ALM neurons) projected onto the choice dimension for catch trials with a 1.8 s delay. Mice were trained with a 1.2 s delay (Chen et al. 2024). Green and orange lines show linear regression fits during the delay and catch periods for right trials, respectively. (d) Distribution over trained networks of the real part of the eigenvalue with the largest real part from the Jacobian matrix, computed at the choice-selective fixed points across 48 trained networks with latent dimensionality *P* = 9–16. Negative values indicate dynamical stability.

Our analysis reveals that the dynamics of the trained networks follows two distinct high-dimensional trajectories encompassing coding and residual dimensions corresponding to each choice. These trajectories transiently align with different dimensions in activity space before settling into selective attractor states corresponding to the two choices. During the delay, the network dynamics evolve along these high-dimensional transient paths, ultimately converging into distinct attractors for left versus right choices (Fig. 3d). Thus, the obtained dynamics reconcile the strong feedforward delay dynamics observed in Daie et al. 2023 with the attractor dynamics observed in ALM in studies where the delay epoch is randomized (Inagaki et al. 2019; Oña-Jodar et al. 2024) (see Discussion).

### Single-trial dynamics

What role do residual dimensions play in shaping individual trials? We incorporated trial-to-trial fluctuations as low-dimensional perturbations of the trial-averaged dynamics (Fig. 5a). We model trial-to-trial variability as trial-dependent changes in low-dimensional input that biases our recurrent network (Fig. 5a,b; see Methods Eq. (26)). The input bias is constant throughout each trial and the recurrent dynamics are constant across trials. Thus, each point in the input bias space (of dimension *P*; Fig. 5d) corresponds to a single-trial trajectory in the activity space (Fig. 5b). Single-trial trajectories are thus represented as perturbations of the trial-averaged low-dimensional flow field (Fig. 5a,b), capturing deviations around the condition-averaged dynamics. These fluctuating inputs may be interpreted as a proxy for state-dependent inputs, e.g., from unobserved regions including neuromodulatory systems, that drift slowly and are not directly tied to task variables. Both the recurrent dynamics and the trial-specific input biases are jointly optimized during training (see Methods).

**Figure 5:**
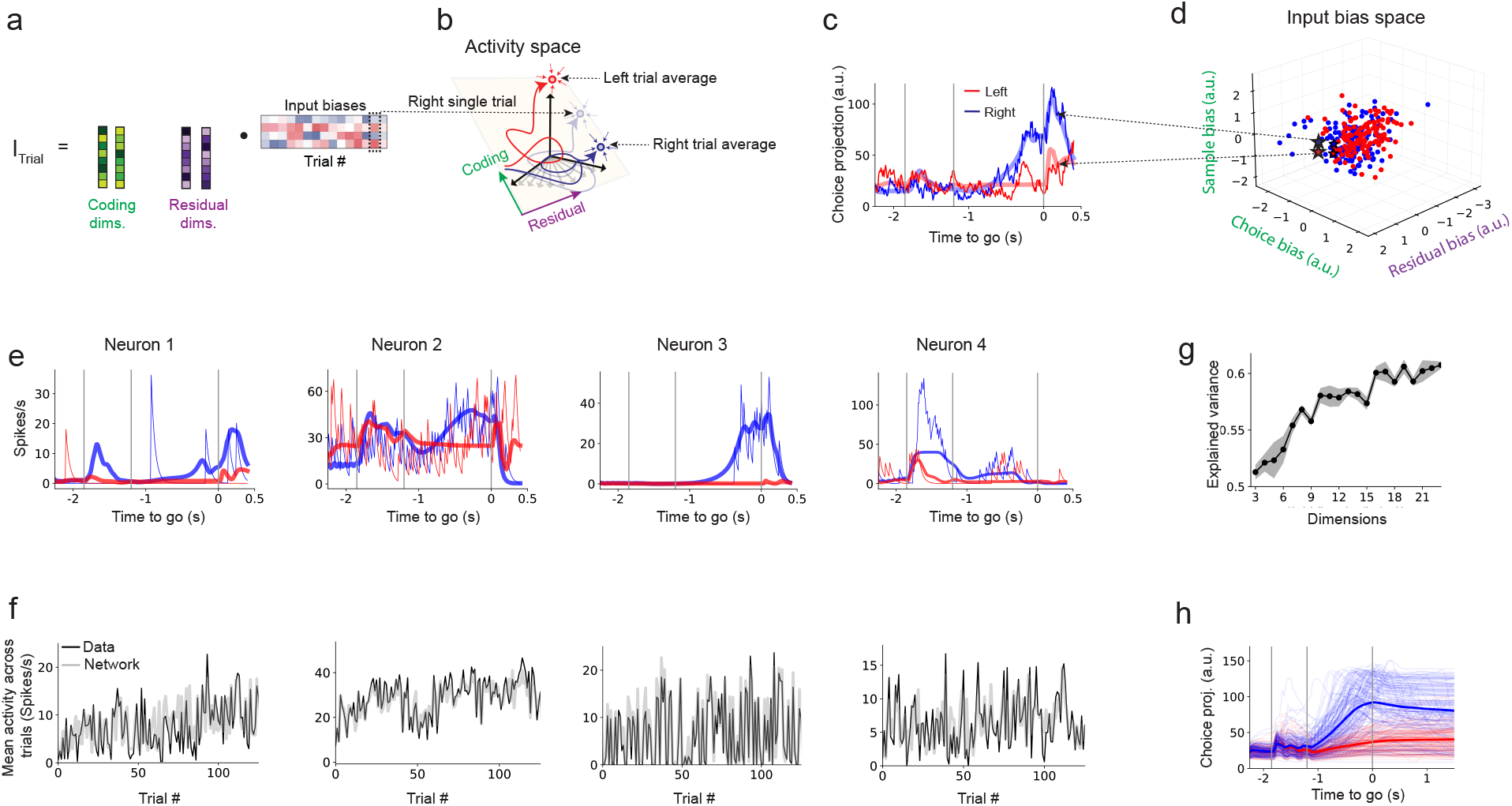
Single-trial dynamics. (a) Low-dimensional, trial-dependent input biases. Biases aligned with coding and residual dimensions are linearly combined to generate input currents that vary across trials. (b) Trial-averaged trajectories for left- and right-correct trials (solid red and blue lines, respectively), and a single right-correct trial (lighter blue line). (c) Single-trial dynamics projected onto the choice dimension for left (red) and right (blue) trials. Solid lines represent recorded data; lighter lines show the corresponding network fits. (d) Three-dimensional projection of the 15-dimensional input bias space, with axes corresponding to the residual, choice, and sample dimensions. Blue and red points correspond to correct right and left trials, respectively. The origin represents the trial-averaged trajectory; deviations from the origin reflect trial-by-trial fluctuations in the input bias. Stars indicate two example single trials shown in panel c. (e) Single-trial activity for four neurons. Thin solid lines: convolved spike trains; thick lines: network fits. (f) Time-averaged activity across trials for the same four example neurons. Black lines: data; gray lines: network fits. (g) Explained variance on the test set across models with different dimensionalities *P* (see Methods). (h) Single-trial network trajectories projected onto the choice dimension for correct left (red) and right (blue) trials when the go cue is withheld. Thin lines: single-trial trajectories; thick lines: condition-averaged trajectory.

We trained the network by simulating multiple trials and constraining it to match several summary statistics of the neural activity in a single session (Fig. S5a). These included: the condition-averaged activity across trials, projections onto coding dimensions (stimulus, choice, response; Fig. 5b; Fig. S5b–c), and time-averages computed within each of the task epochs (sample, delay, response). Training on these summary statistics emphasizes shared structure across neurons and trials (Pellegrino et al. 2024), enabling robust learning of latent dynamics and low-dimensional trial-by-trial variability (see Methods). Input biases span both coding and residual dimensions and modulate single-trial neural dynamics (Fig. 5c; Fig. S5e).

Importantly, the network cannot exploit input biases as a shortcut: withholding the sample stimulus after training causes the network to fail to reproduce single-trial delay activity, confirming that preparatory activity requires stimulus integration and cannot be accounted for by input biases alone (Fig. S5f; see Methods).

Applied to individual recording sessions, the model reliably captured single-trial neural dynamics (Fig. 5c; Fig. S5b–d), including the dynamics of coding dimensions (Fig. S5d), and slow drift in firing rates across trials (Fig. 5f). The model generalized to held-out trials, with cross-validated explained variance saturating at ∼15 dimensions (Fig. 5g). Following training, trial-to-trial bias fluctuations were contained in a subspace spanning both coding and residual dimensions (Fig. 5d; Fig. S5e). Single-trial activity exhibited non-stationary structure over the course of the session, including slow ramping of neural responses across trials (Fig. 5f). This trial-dependent structure is captured by our model but would be missed by approaches that assume independent and identically distributed fluctuations across trials.

With the go cue withheld, the network fit to single trials often exhibits two choice-selective attractors, consistent with the trial-averaged model (Fig. 4; Fig. 5h). The trial-dependent input biases modify the flow field of the network dynamics, altering the local velocity of neural trajectories (Eq. (26)). These changes lead to trial-to-trial variations in the velocity of the choice projection trajectory, as well as shifts in the location and stability of the attractors (Fig. 5h).

Overall, these results demonstrate that the model effectively captures the shared low-dimensional structure across trials.

### Residual dimensions bias decisions on single trials

To study how residual dimensions could bias decisions on single trials, we focused on error trials, instances in which animals made incorrect choices by selecting the lick-port opposite to the stimulus cue. These trials provide a window into the sources of variability that can alter decision outcomes on a single-trial basis. Behavioral errors correlate with trial-by-trial activity fluctuations (Li et al. 2016; Inagaki et al. 2019; Finkelstein et al. 2021). Based on our previous findings using networks trained on condition-averaged activity, we hypothesized that such errors are driven by trial-by-trial fluctuations along residual dimensions, which, while orthogonal to the task-coding subspace, exert a strong influence on the network dynamics.

We reasoned that since trial-by-trial variability in our network arises from fluctuations in input biases along both coding and residual dimensions (Fig. 5d; Fig. S5e), then it should be possible to identify the principal direction of input bias fluctuations that best separates correct and incorrect choices. We refer to this direction as the error axis: the axis in input bias space that most reliably predicts decision errors, which in principle could be different for left and right trials (Fig. 6a). To isolate this axis, we trained linear decoders to distinguish correct from error trials separately for left and right instructed trials, using trial-by-trial input biases extracted from the trained network (Fig. S6a, b; 91% and 93% cross-validated decoder accuracy, respectively).

**Figure 6:**
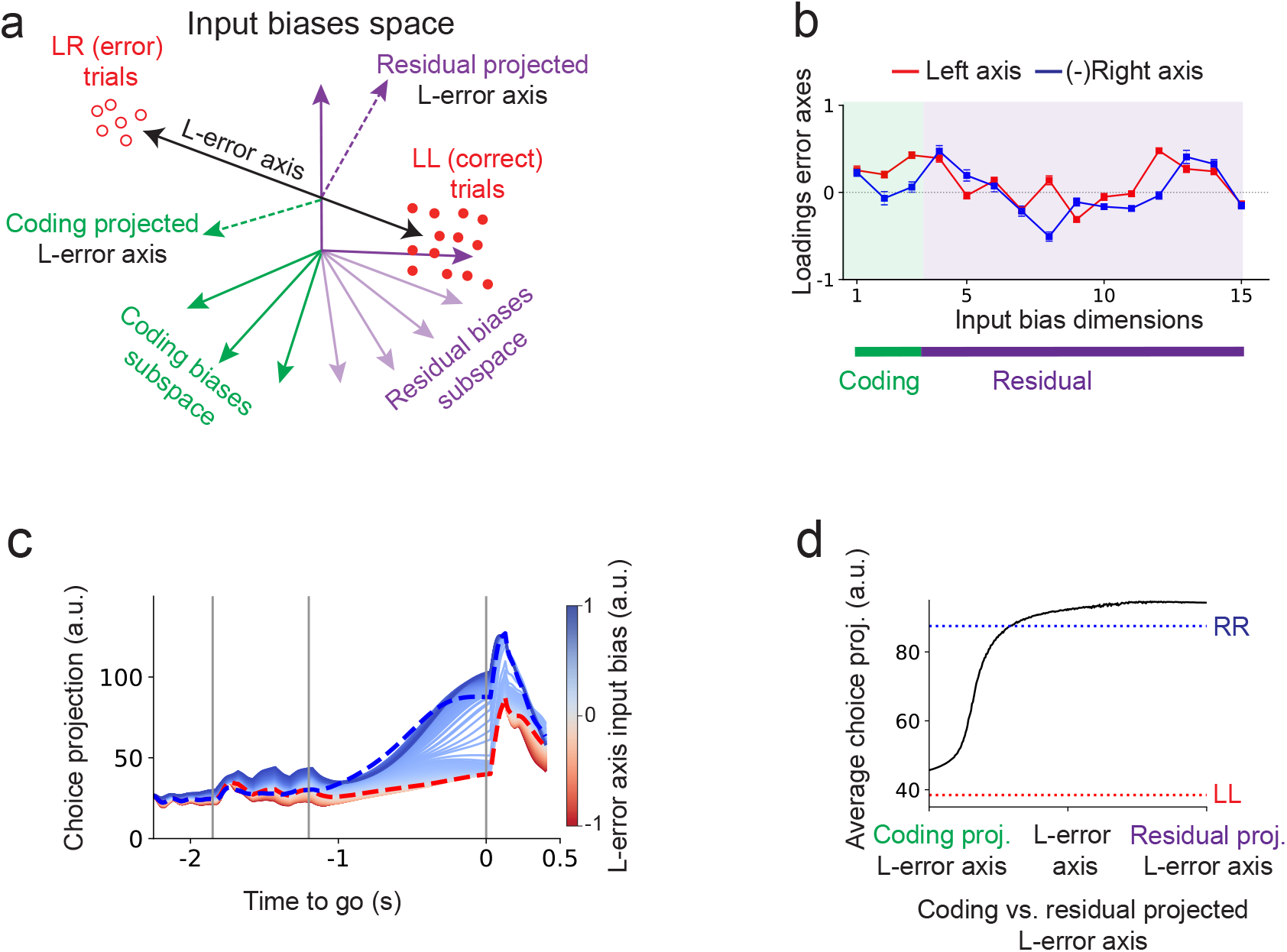
Residual dimensions bias decisions on single trials. (a) The input bias space. Each trial is represented as a point corresponding to the fitted input bias (see Methods). The left error axis (L-error axis) is defined as the direction within the input bias space that differentiates error from correct left trials. The error axis can be decomposed into its projections onto the coding and residual dimensions of the input bias space (coding- and residual-projected L-error axes, respectively). (b) Error axis loadings for classifying correct versus error trials. The first three components correspond to stimulus, choice, and response dimensions, while the remaining components span the residual subspace. Right-trial weights are shown with inverted signs for clarity. Error bars represent the standard deviation computed via bootstrapping (1000 iterations with 80% of trials). Green region corresponds to the coding dimension loadings while the purple region to the residual dimension loadings. (c) Choice projection trajectories for systematic variations of the input bias along the left error axis for left trials. Blue and red dashed lines indicate network’s trial-averaged activity for left and right correct trials, respectively. (d) Mean choice projection during the late delay epoch (215–235 ms). The midpoint of the x-axis corresponds to the input bias being perfectly aligned with the left error axis. The left and right extremes correspond to input biases confined to only coding and only residual components, respectively. Across the x-axis, the composition of the input bias is systematically varied from coding to residual components while its magnitude is held fixed at a value that, when aligned with the left error axis (x-axis midpoint), produces a miss-like left trial (one of the deep blue trials in panel c; see Methods). The underlying choice projection trajectories correspond to those shown in Fig. S6l.

The error axes for left and right classifiers were correlated (Fig. 6b; Pearson correlation between decoder weights: *r* = –0.49), suggesting the presence of a shared component of the left and right error axes that drives decision errors across conditions. Analyzing the error axes loadings revealed contributions from both coding and residual dimensions (Fig. 6b). This suggests that trial-by-trial variability in residual dimensions may contribute to fluctuations in decision outcomes.

To test the causal role of trial-by-trial fluctuations, we systematically varied the network’s input biases along the error axes. These variations in input biases reliably shifted the choice-projected trajectories, converting left trials into right trials and vice versa (Fig. 6c; Fig. S6c-e). Varying the input bias along the error axes gradually shifted the choice projection toward the opposite choice (Fig. 6c; Fig. S6d), consistent with the gradual variations observed across trials (Fig. S6g,j). Despite continuous variation of the input bias along the error axes, the response projection exhibited a sharp shift in trajectories after the go cue, consistent with a discrete-like change in response-evoked activity between left and right correct trials (Fig. S6e, Fig. S6h,k). Inputs along the error axes also induced continuous changes in the sample projection during the sample epoch in the direction of the opposite choice (Fig. S6c), although projections onto the sample dimension alone provided only weak discriminative power between corrects and error trials during this period (Fig. S6f,i).

To isolate the contribution of residual versus coding components to these errors, we constructed input biases of fixed magnitude and systematically varied their composition from fully coding-aligned to fully residual-aligned (Fig. 6a; see Methods). The magnitude was chosen such that, when aligned with the error axis, the input reliably drove choice dimension activity toward values characteristic of error trials (Fig. 6c; Fig. S6d). We then continuously tilted the input bias from the coding toward the residual subspace, isolating whether errors require fluctuations confined to the coding subspace (green region, Fig. 6b), the residual subspace (purple region, Fig. 6b), or a specific mixture of both. Input biases restricted to the coding subspace were insufficient to produce error-like trials. In contrast, input biases with an increasing fraction of residual components reliably redirected choice trajectories toward the opposite choice (Fig. 6d; Fig. S6l–n). These results demonstrate that trial-by-trial fluctuations along residual dimensions are a driver of decision errors.

### Residual dimensions in thalamic activity shape cortical dynamics underlying decisions

Bidirectional coupling between ALM and connected thalamus (parts of the ventromedial, ventrolateral, and medio dorsal nuclei) is causally involved in shaping choice-related cortical neural dynamics in delayed movement tasks (Guo et al. 2017). In our ALM cortical model, residual dimensions play a causal role in shaping decisions. But what role do they play in communication across the thalamocortical loop?

One possibility is that choice-selective neurons in the motor thalamus influence ALM through selective, choice-specific interactions. An alternative scenario is that weakly selective thalamic populations influence cortical dynamics with projections onto residual dimensions. In the latter, thalamic contributions to choice may arise from interactions within residual subspaces. These two possibilities have not been experimentally distinguished (Guo et al. 2017).

We extended our modeling approach to include ALM and connected thalamus (Fig. 7a, Methods). As before, the cortex was modeled as a low-rank recurrent network, capturing both coding and residual dimensions (Fig. 1b). We modeled the thalamus as a non-recurrent population of neurons that is directionally connected with ALM (Guo et al. 2017; Recanatesi et al. 2022). Thalamocortical and corticothalamic long-range connections were modeled as low-dimensional bottlenecks: only a subset of each area’s latent dimensions is transmitted to the other area, reading out along specific combinations of coding and residual dimensions and projecting into a lower-dimensional subspace of the target area (Fig. 7b). This builds on experimental and modeling work suggesting that long-range projections transmit only a subset of the population activity dimensions, typically lower-dimensional than the full latent space. (Semedo et al. 2019; Barbosa et al. 2023; Pereira-Obilinovic et al. 2024; MacDowell et al. 2025). The instructive sensory stimuli were delivered through cortex (Finkelstein et al. 2021), whereas the go cue was routed via thalamus (Inagaki et al. 2022).

**Figure 7:**
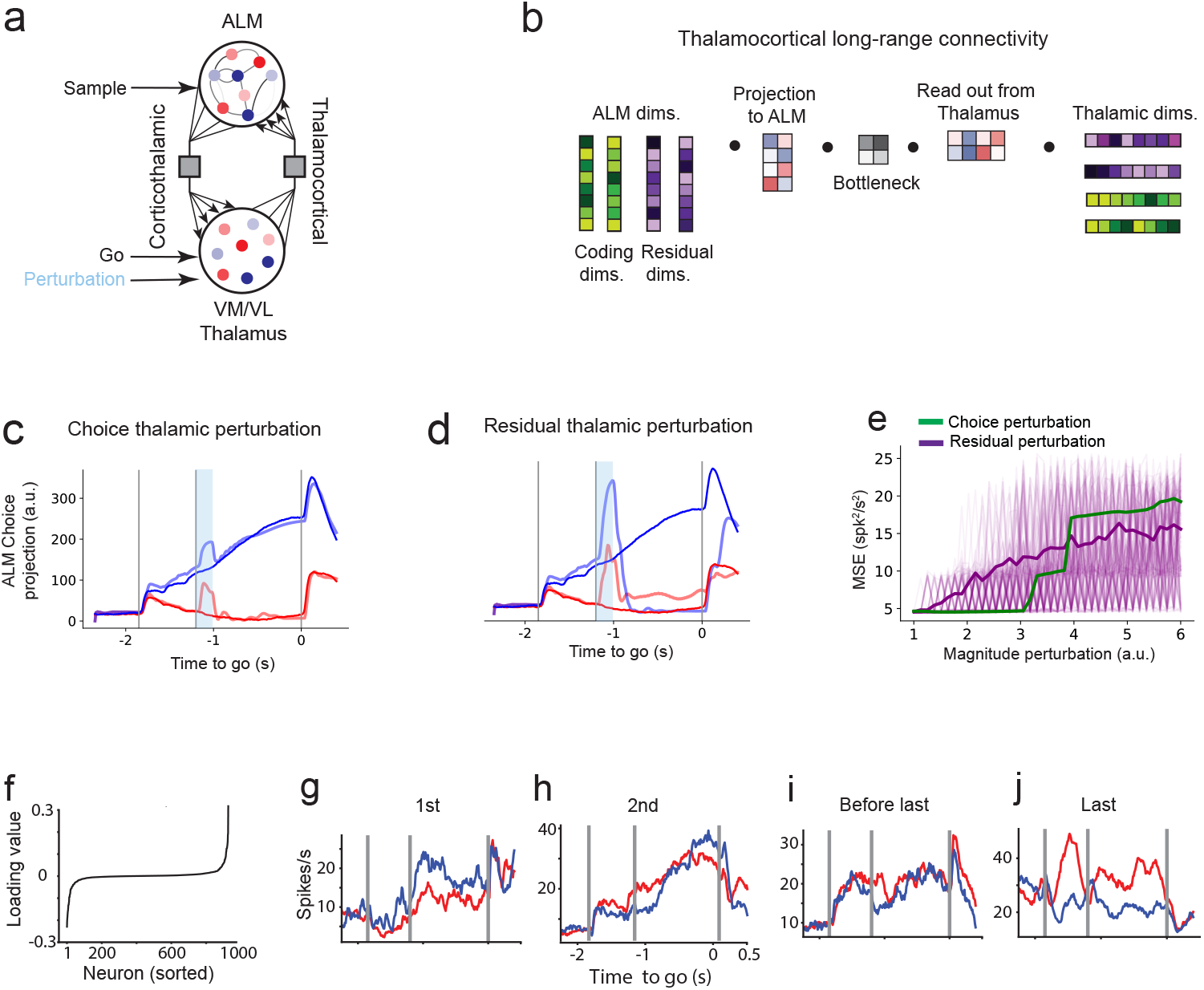
Thalamic residual dimensions shape cortical decision trajectories in a thalamocortical network model. (a) Model including ALM and thalamus. ALM is modeled as a low-rank recurrent network (Fig. 1b); thalamus is a non-recurrent population reciprocally connected to ALM via long-range connections that readout and transmit only a subset of each area’s latent dimensions. Stimuli were delivered through cortex, while the go cue was routed via thalamus. Perturbations are applied in thalamus. (b) Low-dimensional connectivity between ALM and thalamus. (c–d) Choice projection following perturbations in thalamus along the choice dimension (c) and a residual dimension (d). Yellow bar indicates the perturbation period. (e) Mean squared error (MSE) between unperturbed and perturbed trajectories for 120 perturbations along residual (purple) and choice (green) dimensions at increasing magnitudes. Light traces show individual perturbations; solid lines indicate the average. (f) Sorted loadings of thalamic neurons onto the residual dimension shown in (d). (g–j) Trial-averaged activity of neurons with the highest (g–h) and lowest (i–j) loadings from (f) during correct left and right trials.

The network was trained to simultaneously reproduce the trial-averaged activity of ALM and thalamic neurons during correct trials (Fig. S7a, b). Consistent with experimental findings (Guo et al. 2017), we found that disconnecting either the corticothalamic or thalamocortical projections led to a marked reduction in neural activity in the model (Fig. S7c).

Perturbations along the choice dimension in thalamus (Fig.7c, b), despite its strong correlation with behavior, had little effect on choice. In contrast, perturbing a subset of residual dimensions, which have little choice selectivity, consistently disrupt the choice projection (Fig. 7d, e). These effects were robust across hemispheres (Fig. S7d-f).

We examined how individual thalamic neurons contributed to the influential residual dimensions identified in our model. Similarly to the results in ALM (Fig. S2q), neurons with strong loadings onto the influential residual dimensions were largely weakly selective to choice (Fig. 7f-j), and the overall choice selectivity captured by the residual dimension remained weak (Fig. S7g), indicating that the causal influence of these thalamic neurons was not aligned with choice tuning.

Together, these results demonstrate that, within the thalamocortical loop, residual dimensions in thalamus can causally influence decision dynamics. This supports a circuit mechanism in which broadly tuned, weakly selective thalamic populations drive the emergence of cortical selectivity (see Discussion)

## Discussion

We show that neural dynamics in a delayed directional movement task are shaped by neural activity outside task-coding dimensions. We applied a recurrent neural network framework to multi-regional Neuropixels recordings. We find that coding subspaces alone do not autonomously generate decision-related dynamics, but instead depend on interactions with residual dimensions. Although these coding dimensions account for a sizable fraction of the population variance and task selectivity, they depend on interactions with other, weakly selective and residual dimensions. The residual subspaces drive the dynamics, and perturbing them is highly efficient in changing behavioral outcomes. The underlying mechanism is non-normal amplification of neural dynamics along residual dimensions. The neural dynamics occupy a relatively high number of latent dimensions (∼ 10). Weak selectivity along the residual dimensions is dynamically amplified through non-normal interactions in the underlying network. As a result, even small perturbations along residual dimensions can transiently grow and redirect trajectories within the coding subspace. This mechanism explains how seemingly untuned or weakly selective populations can exert strong effects on choice (Daie et al. 2023).

### Dynamical motifs underlying ALM delay activity

Our ALM models reveal a dynamical regime that unifies two dynamical motifs long considered as substrates of short-term memory and decision-making. The network displays strongly non-normal delay dynamics (Goldman 2009; Ganguli et al. 2008; Orhan & Ma 2019; Stroud et al. 2024), yet these trajectories converge onto two choice-selective fixed-point attractor states (Amit & Brunel 1997; Wang 2002; Wong & Wang 2006; Compte et al. 2000; Wimmer et al. 2015). Thus, our data-driven models unify several key observations about ALM dynamics: the strongly non-normal delay activity characterized in targeted perturbation experiments (Daie et al. 2023), the discrete attractor structure demonstrated in random-delay tasks (Inagaki et al. 2019; Oña-Jodar et al. 2024) and in distractor studies (Finkelstein et al. 2021), and the slowdown of neural activity observed in catch trials when the go cue is omitted (Inagaki et al. 2019; Maimon & Assad 2006; Tanaka 2007).

Our results shed new light on the switching dynamics observed in Inagaki et al. (2019). In that study, bilateral photoinhibition occasionally switched population trajectories to the opposite choice. Because broad photoinhibition suppresses activity across all dimensions, these perturbations likely engaged residual dimensions alongside the choice dimension. Our results suggest that the residual component of these perturbations drove the observed switches, consistent with the causal role of residual dimensions identified here.

Our data-driven findings align with the normative theories for how recurrent networks solve delayed-response tasks (Orhan & Ma 2019; Stroud et al. 2023). These studies show that networks trained in delayed-response tasks with fixed delay epochs —in contrast to paradigms with unpredictable delay durations—naturally adopt non-normal dynamics to optimally load sensory information (Stroud et al. 2023): stimulus inputs initially drive activity along amplifying non-normal dimensions, and only later do trajectories collapse onto attractor states during the delay. Our ALM models exhibit these dynamical motifs. Together, our results reveal a unified dynamical view for ALM and connected structures: non-normal transients that funnel activity into choice-selective attractors, with amplification and stability that may be flexibly modulated depending on task demands.

### Residual dimensions shape behavior-related neural dynamics

Classical decision-making models often reduce population activity to a few latent dimensions aligned with stimulus or choice encoding (Wang 2002; Wong & Wang 2006; Mante et al. 2013; Kobak et al. 2016; Inagaki et al. 2019; Langdon & Engel 2025; Luo et al. 2025). These interpretable models largely focus on coding dimensions only and thus overlook important aspects of the population dynamics.

The residual dimensions identified here indicate that neural circuits operate within intermediate-dimensional dynamical subspaces, where additional degrees of freedom interact with coding dimensions to shape decisions. Partial observations can cause data-driven models to spuriously infer line-attractor structure even when the underlying circuit is non-normal (Qian et al. 2024). In our case, the recovered dynamics remain strongly non-normal, suggesting that our model operates at a dimensionality that is in the similar range as that of the ALM delay-epoch dynamics. This view aligns with recent findings in the monkey motor cortex during reaching, where intermediate-dimensional (15-25) non-normal dynamics drive transient activity (O’Shea et al. 2022). Our findings also relate to studies in motor cortex, where preparatory activity unfolds in movement-null subspaces that do not directly drive outputs but modulate execution (Kaufman et al. 2014; Elsayed et al. 2016; Khilkevich et al. 2024). Similarly, we show that residual dimensions can causally shape decision-related dynamics without aligning with canonical choice subspaces.

In the delayed movement task (Fig. 1c) the choice dimension remains relatively stable across the delay epoch (Li et al. 2016; Inagaki et al. 2018; Daie et al. 2023). However, neural activity along the choice dimension is not self-sustaining but instead is driven by dynamics unfolding along both residual and coding dimensions. Choice-selective neurons therefore reflect the upcoming decision but do not control it. These results challenge the conventional view that low-dimensional coding subspaces are sufficient to explain neural computations and reveal that dimensions previously considered task-irrelevant can play a causal role in shaping behavior.

### Modeling trial-to-trial variability as low-dimensional modulation of the global dynamical landscape

In contrast to previous data-driven models that treat trial-by-trial variability as time-varying independent and identically distributed noise (Sourmpis et al. 2023; Pals et al. 2024), we model variability as a non-stationary, low-dimensional modulation of the population dynamics. By representing each trial as a constant input bias acting on both coding and residual dimensions, our framework captures how trial-specific fluctuations perturb the shared low-dimensional dynamics of the network. These perturbations reshape the network’s activity space, effectively reshaping neural trajectories across trials and changing the velocity of the non-normal dynamics, which can be selectively modulated by constant inputs, as recently demonstrated in theoretical models (Gillett & Brunel 2024). This modeling approach is particularly relevant in our task, where contingencies differ but the sensory stimulus within a given contingency is held constant across trials; unlike perceptual decision-making paradigms with trial-varying sensory evidence (Steinemann et al. 2024; Luo et al. 2025), variability here is more likely to reflect fluctuations in internal states (e.g., engagement or satiety) than changes in sensory input. Remarkably, we found that fluctuations along residual dimensions—rather than coding dimensions—were the primary drivers of decision errors, indicating that weakly selective neuronal populations within residual dimensions are causally responsible for fluctuations in single-trial decisions.

### Residual thalamic dimensions drive preparatory dynamics in cortex

Thalamocortical interactions are critical for maintaining preparatory activity in this task (Guo et al. 2017; H. Yang et al. 2022). Photoinhibition of motor thalamus causes a near-complete collapse of ALM activity (Guo et al. 2017), demonstrating its causal role in sustaining cortical dynamics. We considered two possible modes of thalamocortical communication that could generate choice selectivity. The first involves labeled-line projections where choice-selective thalamic neurons connect to matching selective populations in ALM, producing selective delay activity through reverberations within the loop. In this case, perturbations of choice-selective neurons in thalamus would be expected to strongly disrupt selectivity in ALM. The second involves residual projections where weakly selective thalamic neurons, dominated by high loadings in residual dimensions, drive ALM preparatory activity through recurrent thalamocortical interactions. In this case, perturbations along residual dimensions in thalamus would be expected to disrupt selectivity in ALM. Our perturbations in the thalamocortical model show that choice selectivity in the thalamocortical loop arises from residual projections that mediate communication between thalamus and ALM. These results suggest that weakly selective thalamic activity, potentially reflecting inputs from basal ganglia (Wang et al. 2021), may play a key role in controlling thalamocortical dynamics that generate ALM preparatory activity.

### Residual dimensions as control knobs for decision dynamics

From a normative perspective, our findings suggest that neural circuits may use residual dimensions as control knobs to shape decision trajectories. These dimensions could enable fine modulation of specific aspects of the decision process, such as its speed, amplitude, or stability. This view supports a broader reconceptualization of low-variance, weakly task-encoding components of neural activity not as noise, but as organized fluctuations in population activity that reflect flexible neural computations.

## Methods

### Experimental data and preprocessing

#### Neural recordings

We analyzed and trained our models on extracellular recordings from mice performing a delayed response decision-making task (Chen et al. 2024). Recordings were obtained from the anterior lateral motor cortex (ALM) and the ventromedial (VM) and ventrolateral (VL) thalamic nuclei, which are causally involved in this task (Guo et al. 2017). Neural activity was acquired across 179 behavioral sessions from 28 mice, yielding a total of 7,884 cortical neurons (3,640 left hemisphere, 4,244 right hemisphere) and 1,932 thalamic neurons (1,004 left hemisphere, 928 right hemisphere). For the single-trial fits, we used a session with 102 simultaneously recorded ALM neurons in the right hemisphere. Preprocessing steps followed standard filtering and spike sorting pipelines and full data provenance is reported in (Chen et al. 2024). Spikes were then smoothed using an exponential filter with a timescale of 50ms.

#### Identification of task-coding dimensions

To identify the coding subspace, we applied linear discriminant analysis (LDA) to condition-averaged neural trajectories from correct left and right trials (Li et al. 2016; Inagaki et al. 2019; Finkelstein et al. 2021; Chen et al. 2024). This analysis was performed separately for each hemisphere using pseudopopulations constructed by pooling neurons across sessions, resulting in 3,640 neurons for the left ALM and 4,244 neurons for the right ALM. Each trajectory was embedded in the full *N*-dimensional activity space of the corresponding pseudopopulation. The choice dimension identified by LDA was robust to the dimensionality reduction method; demixed PCA (Kobak et al. 2016) yielded similar choice dimensions (cosine similarity equal to 0.87 and 0.85 for the left and right ALM pseudopopulations, respectively).

We extracted the dimensions that best separated left and right trajectories during three key behavioral epochs: (1) the sample dimension 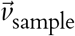, computed from the first 600 ms of the sample epoch; (2) the choice dimension 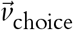, computed from the 600 ms preceding the Go cue; and (3) the response dimension 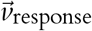, computed from the 350 ms following the Go cue. The resulting vectors 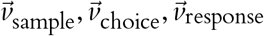 were orthogonalized using the Gram–Schmidt procedure (Golub & Van Loan 2013) to ensure they formed an orthonormal basis for the coding subspace.

These dimensions were selected to reflect the core computational demands of the task: sensory integration (sample), choice maintenance (choice), and motor execution (response). Together, 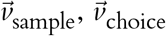, and 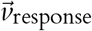 span a coding subspace that captures differences in neural dynamics across conditions at different epochs of the trial. We explicitly incorporated these coding dimensions into the network model through alignment constraints during training (see Training procedure on condition-averaged data).

### Network models

#### ALM model

We modeled ALM as a recurrent rate-based neural network governed by standard rate equations (Grossberg 1969; Hopfield 1984; Miller & Fumarola 2012):

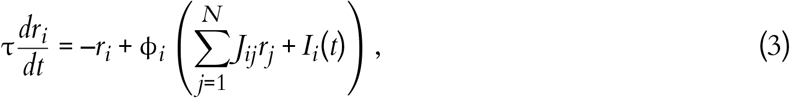

where τ = 40 ms is the neuronal integration time constant. The activation function ϕ_*i*_(*x*) is given by

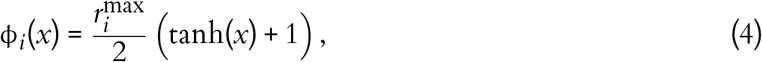

with 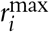 denoting the maximum firing rate of neuron *i*, treated as a trainable parameter. The recurrent connectivity matrix *J*_*ij*_ is constrained to be low-rank and takes the form:

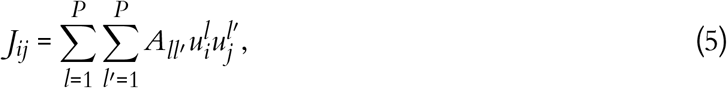

where the vectors 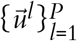 define the connectivity structure. We define the matrix **U** as the matrix whose columns are the coding and residual dimensions, such that 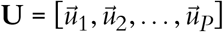. The first *C* vectors span the task-relevant coding subspace, while the remaining *R* = *P*–*C* vectors span the residual subspace. All vectors 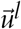 are optimized to remain orthonormal during training (see below). This low-rank structure (Mastrogiuseppe & Ostojic 2018) has been widely adopted in models of associative memory (Hopfield 1982), sequence generation (Sompolinsky & Kanter 1986; Kleinfeld 1986; Gillett et al. 2020), and manifold-based attractor dynamics (Ben-Yishai et al. 1995; Darshan & Rivkind 2022).

The external input is given by:

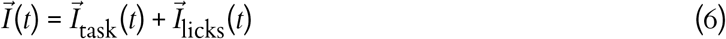

To match the task structure used in the experiments, the external input 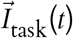 was defined as:

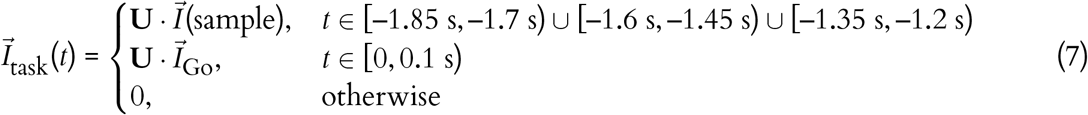

where time zero is the time of the go cue and with sample-dependent input defined as:

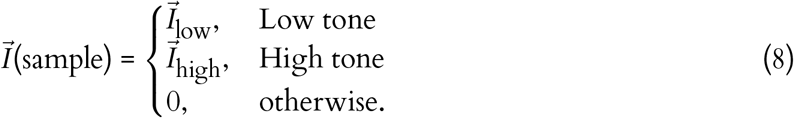

The *P*-dimensional input parameters 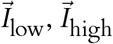, and 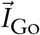 were optimized during training.

Lastly, we included a proprioceptive-like input representing licking activity, which was modeled as a time-varying input proportional to the average lick rate and differed between left and right trials:

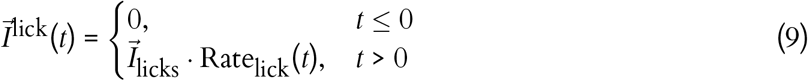

where the amplitude vector 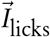 was determined by trial type:

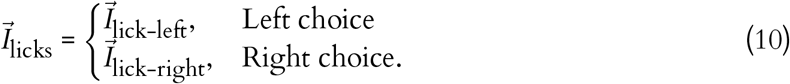

The parameters 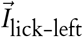 and 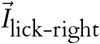 were optimized during training. As indicated by Eq. (9), lick-related inputs only influence the network after the go cue, consistent with the absence of anticipatory licking in behavior. This input captures the strong sensory feedback that licking exerts on ALM, while acknowledging that the motor act of licking originates from downstream structures (Kaku et al. 2025). Thus, we interpret this signal as a proprioceptive input reflecting ongoing motor output. The rationale for including measured licking activity is to prevent the ALM model from having to generate lick-evoked responses that originate outside ALM. However, because the model was trained only up to 350 ms post-go cue, when few licks occur, this input had minimal impact on the network dynamics.

By projecting the network dynamics in Eq. (3) onto the coding and residual subspaces, we obtained a reduced description of population activity in terms of latent variables 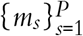, corresponding to the projections of firing rates onto the low-rank basis (Eq. (5)). The resulting latent dynamics evolve as:

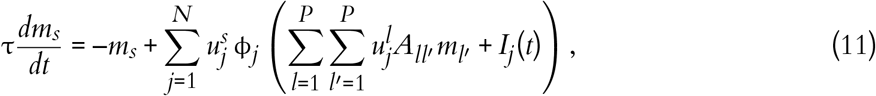

In our implementation, the number of trainable parameters includes *N* × *P* for the low-rank basis vectors 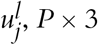 task input vectors, *N* × 2 licking input vectors, *N* maximum firing rates, and *P*^2^ parameters for the interaction matrix **A**.

To assess whether non-normal interactions are necessary to account for ALM dynamics, we trained a network in which the interaction matrix was constrained to be symmetric. Symmetry was enforced by replacing **A** with (**A** + **A**^⊤^)/2 at each forward pass, ensuring symmetry by construction. All other training procedures, hyperparameters, and constraints were identical to the unconstrained network.

For low-dimensional neural dynamics, where *P* ≪ *N*, this low-rank parametrization dramatically reduces the number of trainable parameters to *O*(*N*) compared to full-rank models, which require *O*(*N*^2^) parameters.

### Mathematical analysis of the interaction matrix A

#### Henrici’s departure from normality

To quantify the degree of non-normality in the dynamics matrix **A**, we used the normalized Henrici’s departure from normality (Asllani et al. 2018). This index measures how much a matrix deviates from being normal, that is, from satisfying **AA**^*T*^ = **A**^*T*^**A**, which is equivalent to being unitarily diagonalizable with orthogonal eigenvectors (Golub & Van Loan 2013). The normalized form of Henrici’s measure ranges between 0 and 1.

For a square matrix **A** ∈ ℝ ^*P*×*P*^, it is defined as

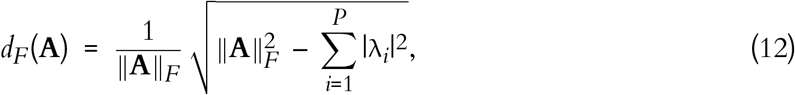

where 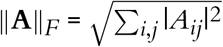 is the Frobenius norm and 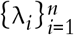 are the eigenvalues of **A**.

By construction, *d*_*F*_(**A**) = 0 if and only if **A** is normal, in which case the eigenvalues fully capture the matrix’s structure. Larger values indicate stronger departures from normality, with *d*_*F*_(**A**) = 1 obtained when all eigenvalues vanish and **A** is nilpotent (e.g. consisting of Jordan blocks with eigenvalue zero). In this extreme case, the spectrum provides no information about the matrix’s behavior, and all the linear dynamics arise from non-orthogonal interactions between eigenvectors (Ganguli et al. 2008; Goldman 2009). While this interpretation is exact for linear systems, in nonlinear networks non-normal interactions can similarly drive strong transient activity (Gillett et al. 2020).

#### Pseudospectra analysis

To quantify the non-normal structure of the linear interactions induced by the interaction matrix **A**, we computed the pseudospectrum. The pseudospectrum characterizes how sensitive a linear system induced by the matrix **A** is to small perturbations and is defined as the set of complex values *z* for which the resolvent norm

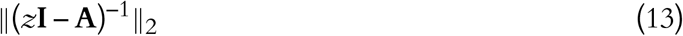

is larger than a threshold (Trefethen & Embree 2020). This norm was computed as the largest singular value of (*z***I** – **A**)^−1^ over a grid of complex values *z* = *x* + *iy* surrounding the eigenvalue spectrum of **A**. Contours of log_10_∥(*z***I** – **A**)^−1^∥_2_ visualize regions where perturbations to coding and residual dimensions are most strongly amplified.

For a normal matrix, whose eigenvectors are orthogonal, the resolvent norm depends solely on the distance between *z* and the eigenvalues,

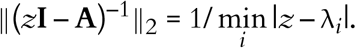

where 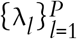 correspond to the eigenvalues of **A**. For a non-normal **A**, where eigenvectors are non-orthogonal, the resolvent norm can become large even far from the spectrum (eigenvalues), revealing directions in latent space where transient amplification occurs. In our model in Eq. (3), the resolvent captures, up to a gain modulation introduced by the network’s nonlinearity, the linearized response of the dynamics to small perturbations in neural activity (see Eq. (18)).

To relate pseudospectral amplification to coding and residual subspaces, we quantified the alignment between the most amplifiable directions and the residual subspace. For each complex *z*, we computed the top right singular vector 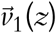 of (*z***I** − **A**)^−1^, corresponding to the input direction that produces maximal amplification. The overlap between 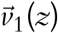 and the residual subspace, spanned by the *P* – *C* dimensional orthonormal basis **R**, was quantified as

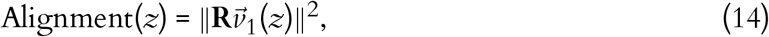

ranging from 0 (fully aligned to the coding dimensions) to 1 (fully aligned to the residual dimensions). This alignment map in Fig. 3h is used to identify which subspaces support maximal transient amplification.

In our fitted models, regions of strongest pseudospectral amplification, corresponding to areas in the complex plane near eigenvalues with positive real parts and with strong shared non-normality among eigenvectors, indicative of non-normal transient amplification, consistently aligned with residual dimensions (Fig. 3h and also b). This analysis shows that transient amplification predominantly acts along these residual dimensions.

#### Schur decomposition and analysis of interaction modes

To characterize the structure of the interaction matrix **A**, we computed its Schur decomposition,

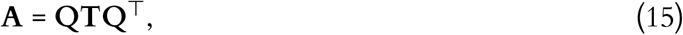

where **Q** is an orthogonal matrix whose columns form an orthonormal basis of activity modes, and **T** is a quasi–lower-triangular matrix whose diagonal entries correspond to the eigenvalues of **A**. In the real Schur decomposition, complex-conjugate eigenvalues appear as 2 × 2 real blocks on the diagonal of **T**, where the off-diagonal element encodes the imaginary part of the eigenvalues and represents the coupling between the corresponding Schur modes. The strictly lower off-diagonal elements of **T** quantify the hidden feedforward structure in the interaction matrix **A** and reflect its degree of non-normality (Hennequin et al. 2012). These terms capture how activity along one Schur dimension can transiently drive or amplify activity in another.

We obtained the Schur-projected activity by projecting the latent variables *m*(*t*) (see Eq. (2)) onto the Schur basis,

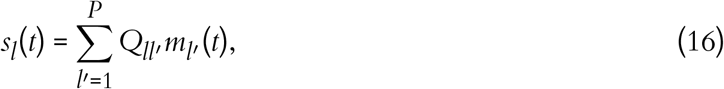

where each component *s*_*l*_(*t*) represents the temporal evolution of the *l*-th Schur mode.

The Schur decomposition is not unique. Within invariant subspaces corresponding to degenerate or complex-conjugate eigenvalues, the columns of **Q** can be freely rotated without changing the decomposition (Golub & Van Loan 2013). In addition, the Schur form allows reordering of the modes, since permuting the diagonal blocks of **T** and the corresponding columns of **Q** preserves **A** = **QTQ**^*T*^. We used this freedom to order the Schur dimensions such that eigenvalues with negative real parts appear first in **T** and those with positive real parts last, ensuring consistent grouping of Schur dimensions across networks.

#### Network perturbations to coding and residual dimensions

To assess the causal contribution of individual network dimensions, we applied brief input perturbations aligned with either coding or residual dimensions during the delay epoch (Fig. 2). Perturbations along the coding and residual dimensions were of the form 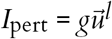, where *g* is the perturbation magnitude and 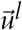 is the *l*-th network dimension. Random perturbations were generated within the subspace orthogonal to the coding and residual dimensions by projecting random vectors onto the column space of **U**^⊥^, whose columns, obtained via QR decomposition of **U**, form an orthonormal basis of its complement.

In Fig. 2g, each perturbation lasted 100 ms and was applied at one of 11 evenly spaced time points during the delay epoch. We tested networks with latent dimensionality ranging from 9 to 16, applying perturbations across all learned dimensions. Each perturbation condition was run on 40 trials, with magnitudes ranging from *g* = –15 to *g* = 15. The Go cue was withheld on both perturbed and unperturbed trials, allowing the network dynamics to evolve toward the corresponding condition-specific fixed point. Simulations ran for 0.9 s following the nominal Go cue time, which in most cases was sufficient for the network to closely approach to the condition-selective attractor states.

To quantify the effect of perturbations, we computed the *rate of choice flips*, defined as the fraction of trials in which the unperturbed activity, when projected on the choice dimension, differed from the choice-projected perturbed activity. Let *x*_L_ and *x*_R_ denote the endpoints of the unperturbed dynamics for left and right trials, respectively. That is, the network output at the end of the delay epoch. Let 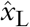 and 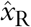 denote the corresponding outputs for perturbed trials. We defined a symmetric threshold margin:

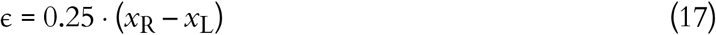

A flip toward right was counted if 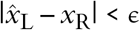, and a flip toward left if 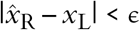. Flip rates were averaged across perturbation trials for each dimension, time point, and magnitude. For Fig. 2g and Fig. S3g-j, we computed the rate of choice flips for 100ms perturbations applied at distinct time windows (during the delay for Fig. 2h; during the sample and delay epochs for Fig. S3g-j) across network dimensions. For each dimension, the perturbation amplitude was varied from –15 to 15. In Fig. 2h, perturbations were aligned with the choice and residual dimensions; in Fig. S3g-j, they aligned with the Schur dimensions (see Mathematical analysis of the interaction matrix).

#### Network perturbations targeting choice- and residual-selective neurons

Current experimentally feasible manipulations act on subsets of neurons rather than specific dimensions in latent spaces (Daie et al. 2021). To mirror this constraint, we targeted neurons most aligned with either the choice or residual subspaces, instead of perturbing the macroscopic dimensions themselves.

Using the decoder matrix **U** (Eq. (5)), we computed per-neuron selectivity with respect to the choice and residual dimensions by sorting the loadings of these dimensions. For a given group size *K* ∈ {1, …, 30}, we selected the top-*K* neurons by the chosen ranking (“choice-selective” or “residual-selective”).

Perturbations were implemented as an external input 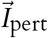 delivered during the first 100 ms of the delay epoch (see Fig. 2j-l). For a target class (choice or residual), group size *K*, and magnitude α ∈ {10, 20, 30}, each neuron in the selected set received input of amplitude 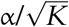. Thus, 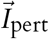 is *K*-hot with constant norm across *K*: 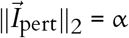.

To quantify the effect (Fig. 2j), we computed the absolute difference between the left- and right-trial trajectories projected onto the choice axis, averaged over the 100 ms preceding the Go cue, for perturbations to choice-selective and residual-selective subsets.

#### Stability analysis of fixed-point attractors

To assess the stability of condition-specific fixed points reached during the delay epoch, we analyzed the linearized dynamics of the trained ALM network models using the Jacobian evaluated at the final state of the trial-averaged trajectories.

We evaluated models trained on right hemisphere ALM activity, with latent dimensionalities ranging from *P* = 9 to *P* = 16, including multiple random initializations per dimensionality. For each model, we simulated the network response to trial-averaged inputs corresponding to left and right conditions, withholding the Go cue, and allowed the network to evolve toward a fixed point. To ensure convergence, we extracted the final network state at time 1.05s after the Go cue.

The Jacobian was computed by linearizing the dynamics in Eq. (3) around the final state. The linearized system governing small perturbations 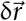 near the fixed point takes the form:

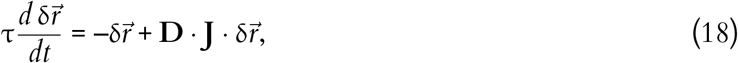

where **D** is a diagonal matrix of derivatives of the activation function ϕ_*i*_ (Eq. (4)), and **J** is the network connectivity matrix:

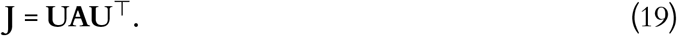

The gain-modulated activation derivative for each neuron is given by:

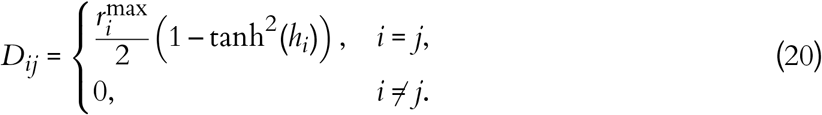

where *h*_*i*_ = ∑_*j*_ *J*_*ij*_*r*_*j*_ + *I*_*i*_(*t*) is the input current to neuron *i* at the fixed point.

The Jacobian matrix, **M** = –*I* + **DJ**, was constructed using Eqs. (19,20) and used to compute the eigenvalues separately for left and right conditions. The eigenvalue with the largest real part determined the local stability of the fixed point (Guckenheimer & Holmes 2013): negative real parts indicate stability, while values near zero or positive indicate marginal or unstable dynamics.

This analysis was repeated across all trained models, yielding a distribution of maximal eigenvalues as a function of latent dimensionality and random seed (Fig. 4d, which corresponds to 48 trained networks), quantifying the stability of task-specific attractor states in our low-rank models of ALM.

#### Analysis of catch trials

In a subset of sessions from the dataset in Chen et al. 2024, the go cue was delayed from 1.2 s to 1.8 s on a small fraction of trials. We refer to these as catch trials. Catch trials were present in 101 out of 179 sessions, comprising a total of 2,529 recorded neurons in right ALM.

To test whether population dynamics stabilize after reaching a choice-selective attractor during catch trials, we quantified the slope of activity projected onto the choice dimension across two periods within the delay epoch: the original delay period (0–1.2 s) and the extended delay period corresponding to the catch interval (1.2–1.8 s). Slopes were computed from the trial-averaged choice projection for right-choice trials in each period.

To quantify uncertainty in slope estimates, we performed a neuron-subsampling bootstrap. On each bootstrap iteration, 1,500 neurons were randomly subsampled without replacement, the choice dimension was recomputed from the subsampled population, and neural activity was projected onto this dimension. Linear slopes were then fit separately in the original and extended delay periods. This procedure was repeated 30,000 times to obtain bootstrap distributions of slope estimates.

From these distributions, we computed confidence intervals and tested two hypotheses: (i) that the slope during the original delay period was positive, consistent with ramping along the choice dimension, and (ii) that the slope during the extended delay period was not significantly different from zero, consistent with stabilization of the dynamics after reaching a choice-selective attractor.

To quantify whether the slope of choice-projected activity differed between the two periods, we computed on each bootstrap iteration the difference between the slope during the original delay period and the slope during the catch interval. We tested the one-sided hypothesis that this difference was greater than zero, consistent with a transition from ramping dynamics during the original delay to stabilization during the catch interval, with *p* defined as the fraction of bootstrap iterations in which the difference was less than or equal to zero.

#### Training procedure on condition-averaged data

We trained the network model described in Eqs. (3–11) to reproduce trial-averaged neural dynamics recorded in the ALM during a memory-guided decision-making task only for correct trials. The training objective minimized a weighted sum of three loss terms:

1. **Reconstruction loss**. This term minimized the mean squared error between the model’s predicted firing rates, 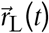 and 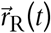, and the condition-averaged neural data, 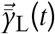 and 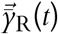, for left and right choice trials:

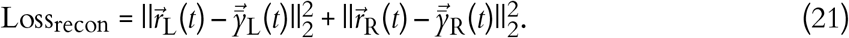
2. **Coding alignment loss**. The model was trained to align its coding dimensions 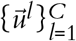 to match the axes extracted from the neural recordings that optimally encode sample, choice, and response using LDA 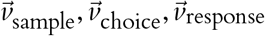. For this we introduced a cosine similarity loss:

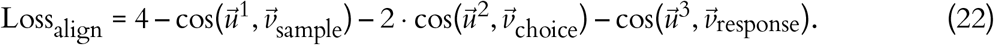

The cosine similarity term for the choice axis was weighted more heavily (multiplied by 2) to prioritize alignment with the choice dimension, which serves as the network’s primary readout for upcoming decisions.
3. **Orthogonality loss**. To enforce orthogonality among the low-rank connectivity dimensions, we penalized deviations of the decoder matrix *U* from an orthonormal basis:

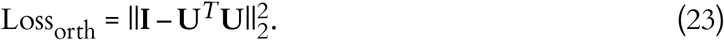

The total loss combined these terms:

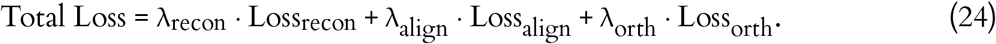

Loss weights were set to λ_recon_ = 0.7, λ_align_ = 0.5, and λ_orth_ = 500. These values were chosen based on empirical performance across fits.

Training was performed using the Adam optimizer with a learning rate of 5 × 10^−4^ for up to 400,000 iterations. Gradients were clipped to improve training stability. Separate models were trained for each hemisphere using custom scripts in PyTorch.

All models were implemented in PyTorch 2.4.0 with CUDA 12.4.0 and trained on an NVIDIA T4 GPU. Each model converged in under approximately 36 hours. We trained multiple models per hemisphere across a range of latent dimensionalities (*P* = 3–20) and at least five random seeds per setting.

#### Cross-validation for networks trained on trial-averaged neural recordings

To assess generalization, we performed cross-validation by repeatedly splitting the dataset at the trial level. For each fold, 80% of the trials were randomly sampled to compute condition-averaged neural activity and train the network. The remaining 20% were held out to evaluate model performance.

This procedure was repeated across five independent random splits using different seeds. For each fold, we computed the mean squared error (MSE) between the network’s predicted firing rates and the trial-averaged activity in the held-out test set. We also computed the cosine similarity between the coding dimensions extracted from the network and those derived from the data using LDA, both computed on the training set. All reported metrics reflect the average and standard error across folds (Figs. 1 and S1).

This procedure was repeated across a range of latent dimensionalities and separately for each hemisphere (Fig S1) and averaged over hemispheres (Fig 1). The results were smoothed using a five-point moving average.

To quantify how much of the condition-dependent neural activity is captured by each coding and residual dimension, we computed the fraction of trial-average variance explained following (Inagaki et al. 2018). For each task epoch (sample, delay, response), we averaged the condition-averaged selectivity (right minus left correct trial-averaged activity) over time, yielding a population selectivity vector of length *N*. The fraction of variance explained by each dimension was then computed as the ratio of the squared projection of this vector onto that dimension to the total variance of the population selectivity vector, defined as the sum of squared entries across neurons. To evaluate the contribution of each dimension to model performance, we computed explained variance on the concatenated trial-averaged activity across both conditions (W. Yang et al. 2022). These two measures serve distinct purposes: the first quantifies how much of the condition-dependent variance is captured by each subspace, while the second evaluates overall model performance by assessing how accurately the network reproduces the trial-averaged firing rates of each condition.

#### Model of single-trial variability

To capture trial-by-trial variability, we modeled each trial as receiving a low-dimensional, trial-specific input current. Specifically, on trial *k*, the input to neuron *i* was defined as:

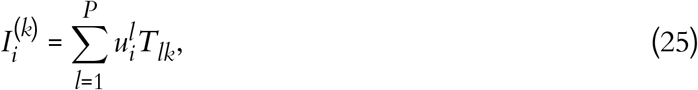

where *T*_*lk*_ is a trial-specific matrix of coefficients, and 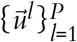 are the low-rank basis vectors spanning the coding and residual subspaces (Eq. (5)). These inputs define a constant offset along the low-dimensional subspace for each trial, shifting the network’s operating point without modifying its connectivity.

The rate dynamics *r*^(*k*)^ for neuron *i* on trial *k* were then governed by:

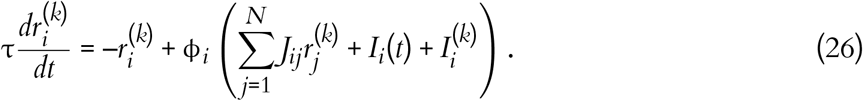

The inputs 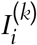 remained fixed throughout each trial but varied across trials. This formulation introduces variability across trials without modifying the recurrent connectivity.

These inputs can be interpreted as introducing small, trial-specific deformations in the vector field governing the dynamics in Eq. (3). Because the variability is restricted to the coding and residual subspaces, it remains interpretable and aligned with the dominant dimensions that shape population activity.

This approach was motivated by the non-stationary fluctuations observed in neural recordings (Fig. 5f), which likely reflect changes in internal states such as arousal, engagement, or satiety, which are factors known to modulate cortical activity (Poulet & Petersen 2008; Matteucci et al. 2022; Steinmetz et al. 2019; Allen et al. 2019). Unlike additive independent and identically distributed noise models, this formulation captures trial-to-trial variability as a low-dimensional, structured modulation of the network dynamics.

#### Training procedure on single-trial data

We trained the single-trial model described in Eq. (26) to capture structured, low-dimensional variability across trials. The single-trial neural data were represented as a tensor *y*_*kti*_, where *k* indexes trials, *t* time points, and *i* neurons. Training was performed by jointly optimizing the parameters of the recurrent network (Eqs. (3–11)) and the trial-specific input currents 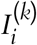 (Eq. (25)) to reproduce summary statistics of this tensor. These trial-specific inputs perturb the network along low-dimensional directions within the coding and residual subspaces (Eq. 5), enabling the model to capture trial-to-trial variability in an interpretable manner aligned with the network’s low-dimensional structure.

In addition to the low-dimensional, constant trial-specific input currents, the model also received the condition-averaged task inputs described in Eqs.(6–10), as well as convolved single-trial lick signals. These lick inputs were weighted separately for left and right responses using the trained input weights defined in Eqs.(9, 10). They were introduced to match lick-related activity likely originating from subcortical structures (Kaku et al. 2025) and influenced the network dynamics only after the Go cue, since animals did not lick during the sample or delay epochs. As in the trial-averaged network, including measured licking activity prevents the ALM model from having to generate lick-evoked responses internally; omitting these inputs would otherwise force the model to produce activity that originates outside ALM.

To prevent the model from learning a direct mapping from the trial-specific input current 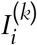 to the responses, we jittered the trial start time at each training epoch. On each trial *k*, inputs began at an offset independent and identically uniformly distributed across trials 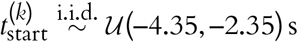 relative to the go cue. Training networks with a fixed start time, they typically exploited input biases to produce transient activity that fits the data without integrating stimulus information to generate delay activity. Jittering the start time during training prevents the network from finding this solution and forces it to integrate stimulus information over time rather than relying on 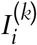 alone. After training, removing the stimulus inputs caused the trained networks to fail to reproduce the variable ramping of the choice latent characteristic of single-trial neural activity, indicating that integration of sample information is required to generate preparatory activity (Fig. S5m).

Rather than fitting individual spike trains, we trained the model to reproduce multiple contractions of the neural activity tensor *y*_*kti*_. This approach leverages the strengths of rate-based modeling while capturing critical statistical features of trial-to-trial variability inherent in spiking data. Our primary goal was to uncover low-dimensional neural structures that generalize across trials. Direct fitting to individual spike trains risks overfitting to trial-specific noise, limiting the model’s generalization. In contrast, training the model using summary statistics (tensor contractions) emphasizes shared structure across trials, promoting robust learning of stable latent dynamics and interpretable low-dimensional variability (Pellegrino et al. 2024) (Fig. 5). Additionally, we explicitly included contractions aimed at capturing variability within the coding dimensions. This allowed us to investigate mechanisms underlying trial-to-trial variability specifically within task-relevant dimensions, such as the choice axis, providing insights particularly relevant to the analysis of error trials. The following losses were minimized:

1. **Trial-averaged loss**. Matches the model firing rates to the condition-averaged neural activity across trials, computed separately for left and right choice conditions as in Eq. (21).
2. **Epoch-averaged loss**. Matches average firing rates across trials and neurons within task-defined temporal epochs:

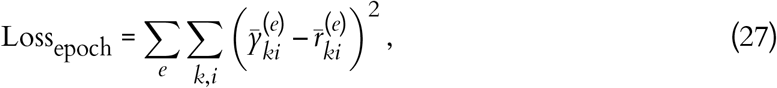

where 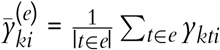 and 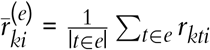 are the trial- and epoch-averaged firing rates for neuron *i* on trial *k* during epoch *e* ∈ {sample, delay, response}.
3. **Projection loss**. Matches the model’s projections onto the coding dimensions 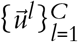 to the single-trial neural activity projected onto LDA-computed coding dimensions:

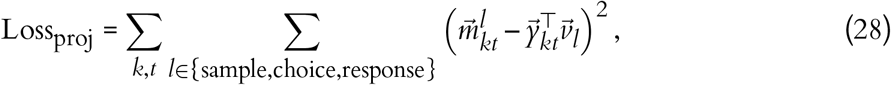

where 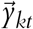 is the observed population activity on trial *k* at time *t*, and 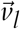 is the LDA-derived coding axis for dimension *l* ∈ {sample, choice, response}. The model projections 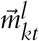 are computed as the dot product between the model’s firing rates and the corresponding coding direction: 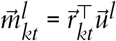. The coding dimensions 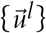 and the LDA vectors 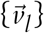 are encouraged to be aligned via the coding alignment loss (Eq. (22)) but are not necessarily identical. This loss encourages the model to reproduce trial-by-trial variability along coding dimensions and further reinforces alignment between the model’s learned coding subspace and the LDA-derived axes.
4. **Coding alignment loss**. See Eq. (22).
5. **Orthogonality loss**. See Eq. (23).
6. **Input regularization**. This term penalized the overall magnitude of the trial-specific inputs to constrain their influence on the dynamics:

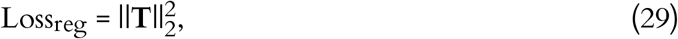

where **T** is a matrix with one column per trial and one row per low-dimensional input direction (Eq. (25)). This constraint encourages the model to account for trial-by-trial variability through modest perturbations, ensuring that most structure arises from the low-dimensional dynamics.

The total loss was a weighted sum of all six components:

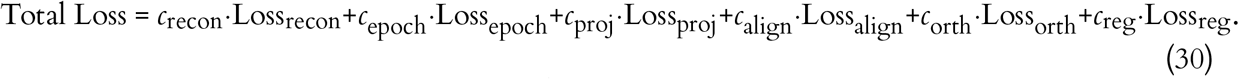

Loss weights were selected empirically (manual tuning to promote stable optimization and a consistent decrease in the loss). We used *c*_recon_ = 0.65, *c*_epoch_ = 0.1666, *c*_proj_ = 0.02, *c*_align_ = 0.5, *c*_orth_ = 500, and *c*_reg_ = 0.1. To stabilize training, full loss terms were activated only after the trial-averaged loss fell below a threshold.

Training used the Adam optimizer with learning rate 5 × 10^−4^ and gradient clipping. All models were implemented in PyTorch and trained on NVIDIA T4 GPUs, with convergence typically reached in under 72 hours.

#### Single-trial cross-validation procedure

After training the model on population-level statistics as described in the previous section, we assessed generalization by splitting the correct trials into a training set (80%) and a test set (20%). The model was trained exclusively on the training set. At test time, only the trial-specific input currents 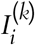 were optimized, while all other model parameters were held fixed. All reported analyses were performed on the held-out test trials.

The rationale behind this cross-validation procedure is that the network, through its low-rank recurrent connectivity (Eq. (26)), learns low-dimensional dynamics that reflect structure that is shared across trials. Therefore, when perturbed through trial-specific low-dimensional inputs, the network can reproduce individual test trials without modifying its recurrent connectivity. This demonstrates that the learned dynamics generalize across trials through low-dimensional perturbations of the network’s operating point.

#### Error axis estimation from input biases

To test whether trial-by-trial variability in residual dimensions could bias decisions, we focused on error trials, where animals selected the incorrect lick port. For each trial, we extracted the trial-specific input biases (Eq. (25)), yielding a single feature vector in the input bias space. Correct and error trials were then labeled, and linear classifiers (logistic regression with no intercept) were trained separately for left and right choices to distinguish correct from error trials. Performance was assessed using an 80/20 stratified split, and standard errors were estimated by bootstrap resampling (1000 iterations).

The normalized decoder weights define the error axes for left 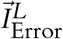 and 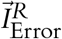 trials, the direction in the input bias space that best separates correct from error trials. Unlike coding and residual dimensions, which are defined in the neuronal activity space of dimension *N*, the error axis lies in the lower-dimensional input bias space *P*.

We systematically generated error trials by adding input biases to the network (Eq. (25)) along the left and right error axes.

#### Perturbations along coding and residual components of the error axes

To investigate the contribution of coding and residual dimensions to error generation, we provided the network with input biases that systematically varied in their alignment with the empirically defined error axes 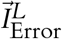 and 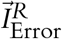 ranging from purely residual to purely coding alignment. The total magnitude of each input bias was kept constant at a value sufficient to generate error-like trajectories when applied along the unmodified error axis.

To dissociate the relative contribution of coding and residual components, we continuously varied the composition of the input biases while keeping their total magnitude constant. Each input bias was constructed by linearly combining the coding and residual components of the error axis,

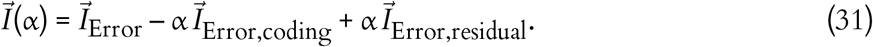

This interpolation spans from purely coding (α = –1) to purely residual (α = 1) perturbations. Each resulting input bias was normalized and applied to the network with the same overall magnitude, generating families of single-trial trajectories. These trajectories were then used to quantify how the coding and residual components of the left and right error axes contribute to the emergence of error-like single-trial neural dynamics (Fig. 6c and Fig. S6c–e).

### Model of ALM–Ventral Thalamus in a Memory-Guided Decision-Making Task

#### Thalamocortical loop

We extended the ALM model (Eqs. (3–11)) to include a reciprocal loop with ventral thalamus. ALM and thalamus were modeled as interacting low-rank networks coupled through dimensionality bottlenecks (see below). Based on anatomical evidence, thalamus was treated as a feedforward population: it receives cortical input via corticothalamic projections and sends output back to ALM via thalamocortical projections, without recurrent connections. The dynamics were:

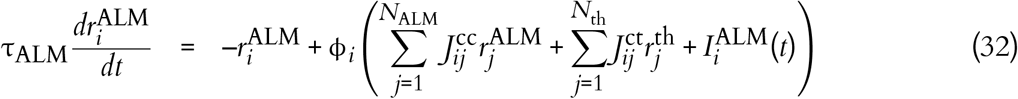

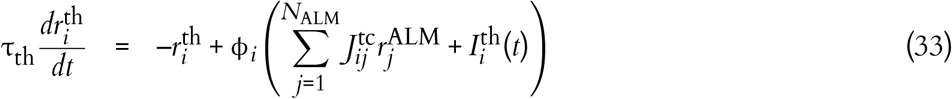

Here, **J**^cc^ denotes recurrent cortical connectivity, **J**^ct^ the thalamocortical projections, and **J**^tc^ the corticothalamic projections. In line with experimental evidence, the thalamic integration time constant was set to one third of the cortical value (τ^th^ = τ^ALM^/3 = 40/3 ms). Consistent with anatomical data, no recurrent connectivity was included within thalamus.

#### ALM local connectivity

ALM connectivity was written similarly as in Eq. (5),

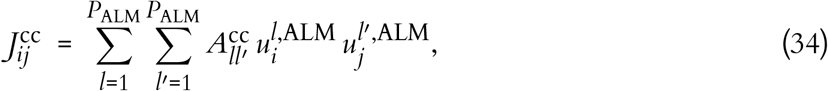

with **A**^cc^ trainable.

#### Thalamocortical and corticothalamic connectivity

We modeled thalamocortical and corticothalamic projections as operating through low-dimensional bottlenecks in latent space (Barbosa et al. 2023; Clark & Beiran 2024; Pereira-Obilinovic et al. 2024). This reflects the idea that only a restricted set of activity patterns is transmitted between ALM and thalamus, while the remaining dimensions remain local, as shown previously for cortico-cortical communication (Semedo et al. 2019; MacDowell et al. 2025).

In the model, each projection was written as a product of three matrices: a readout from the presynaptic subspace, a bottleneck that mixes signals in a reduced set of dimensions, and a projection into the postsynaptic subspace (see Fig. 7b):

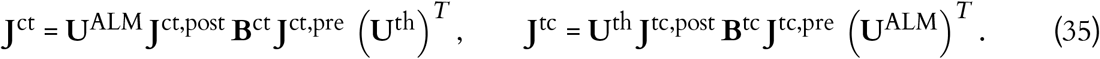

Here **U**^ALM^ and **U**^th^ span the coding and residual subspaces in ALM (see Eq. (34)) and thalamus, respectively. For the thalamocortical projections, **J**^ct,pre^ selects a *P*^ct^-dimensional subspace of the thalamic latent space (*P*^ct^ ≤ *P*^th^). These signals are mixed within the bottleneck **B**^ct^ and projected into a *P*^ct^-dimensional subspace of ALM defined by **J**^ct,post^ (*P*^ct^ ≤ *P*^ALM^).

Analogously, corticothalamic communication is restricted to a *P*^tc^-dimensional subspace of ALM selected by **J**^tc,pre^, transformed through **B**^tc^, and projected into thalamus by **J**^tc,post^.

Thus, despite the higher dimensionality of both ALM and thalamus, reciprocal communication is limited to two narrow subspaces: *P*^ct^ thalamic dimensions transmitted to ALM and *P*^tc^ ALM dimensions transmitted to thalamus. The remaining dimensions in each area remain private and do not propagate across the loop.

For Fig. 7, we set the latent dimensionality of ALM and thalamus to *P*^ALM^ = *P*^th^ = 10, with both the thalamocortical and corticothalamic bottlenecks restricted to *P*^ct^ = *P*^tc^ = 5.

Lastly, the external inputs to the network followed the same structure as described in Eqs.(6–10), with one key difference: consistent with experimental findings (Inagaki et al. 2022), the go cue was delivered to the thalamic network, whereas the sample stimulus was delivered to ALM.

#### Training procedure

Training followed the same procedure described for the cortical model described previously, but was extended to include both ALM and thalamus. The reconstruction loss was computed jointly across the two areas, requiring the model to reproduce condition-averaged activity in both ALM and thalamus. Coding alignment losses were applied separately in each area, constraining the learned low-rank subspaces (**U**^ALM^, **U**^th^) so that their coding dimensions aligned with task-relevant axes extracted from ALM or thalamic recordings. Orthogonality losses were likewise imposed independently within ALM and thalamus. The total loss was a weighted combination of reconstruction, alignment, and orthogonality terms, using the same coefficients as in the cortical model. Models were trained separately for left and right hemispheres with the Adam optimizer (learning rate 5 × 10^−4^) and gradient clipping.

#### Random perturbations in thalamus

In Fig. 7e, following the perturbation analysis described above (see Network perturbations), we tested the sensitivity of the thalamocortical model to random inputs applied to thalamus. Perturbations were of the form

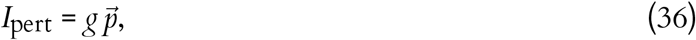

where *g* is the perturbation magnitude and 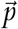 is either the thalamic choice axis or a random direction within the residual subspace. For 40 magnitudes (linearly spaced between 1 and 6), we applied inputs along the choice axis or along 120 random residual directions. Random vectors were sampled from a standard normal distribution and normalized to unit norm.

The effect of each perturbation was quantified as the mean squared reconstruction error between perturbed model activity and condition-averaged neural activity in ALM and thalamus. Losses were compared across choice-axis and residual-subspace perturbations to assess the relative influence of task-relevant versus residual dimensions.

## Data and code availability

All analysis and modeling code supporting this study is distributed across six GitHub repositories:

1. Training the network models on condition-averaged data: https://github.com/ulisespereira/alm-coding-residuals-condavg-train
2. Analysis of the network models trained on condition-averaged data: https://github.com/ulisespereira/alm-coding-residuals-condavg-analysis
3. Training the network models on single-trial data: https://github.com/ulisespereira/alm-coding-residuals-singletrial-train
4. Analysis of the network models trained on single-trial data: https://github.com/ulisespereira/alm-coding-residuals-singletrial-analysis
5. Training the thalamocortical network models on condition-averaged data: https://github.com/ulisespereira/alm-thal-coding-residuals-condavg-train
6. Analysis of the thalamocortical network models trained on condition-averaged data: https://github.com/ulisespereira/alm-thal-coding-residuals-condavg-analysis

Together, these repositories provide the complete modeling pipeline required to reproduce the results reported in this study.

## Acknowledgments

We would like to thank Nicolas Brunel, Jeremiah Cohen, Kabir Dabholkar, Laura Driscoll, Arseny Finkelstein, David Hansel, Shani Kaminitz, Shawn Olsen, Alessandro Sanzeni, and Joshua Siegle for useful comments and discussions.

This work was supported by Allen Institute, National Institute of Neurological Disorders and Stroke (NINDS), National Institute on Drug Abuse (NIDA) of the National Institutes of Health under Award Number U19NS123714 and Israel Science Foundation (ISF, 2097/24 and 2994/24). R.D. was supported by the Alon scholarship for the integration of outstanding faculty.

## Author contribution matrix

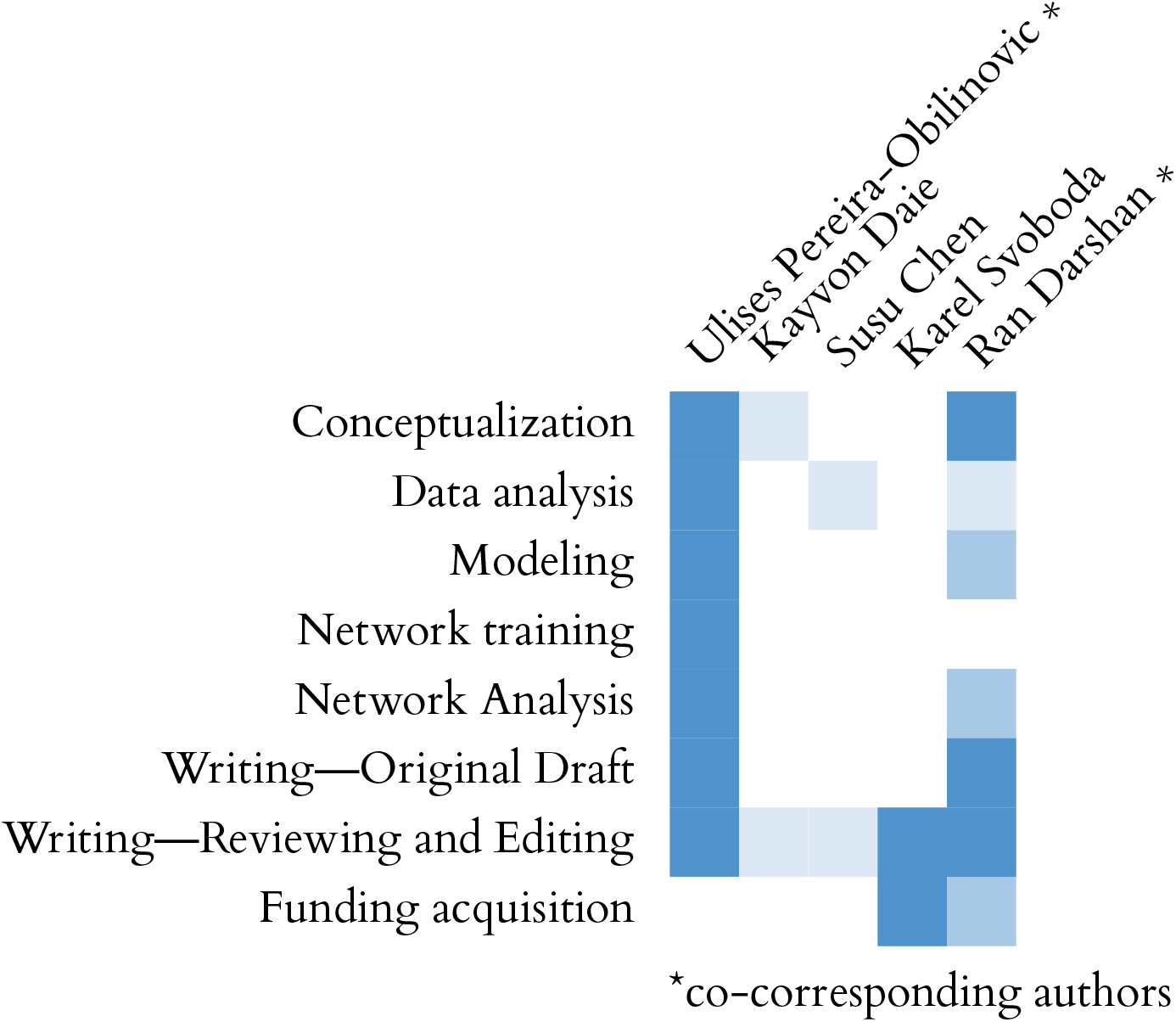

---

**Figure S1:**
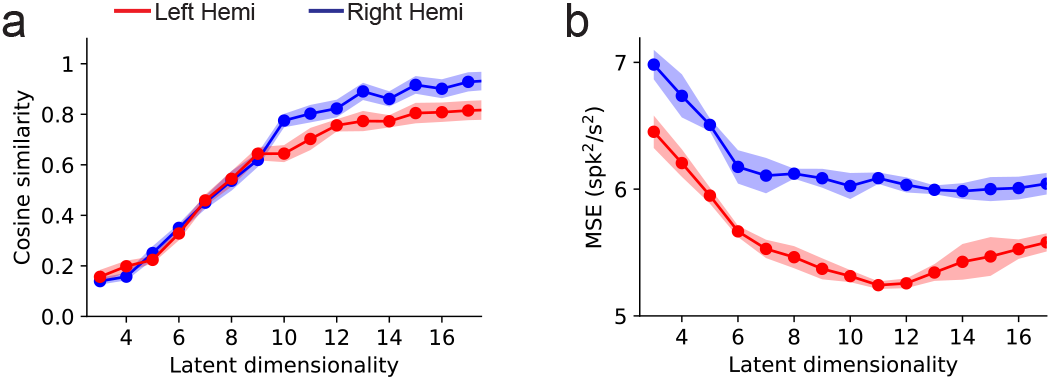
Supplementary Figure corresponding to Fig. 1. (a) Average cosine similarity between the coding dimensions extracted from the network fit and those derived from the data, computed across five independent subsets using 80% of the trials to estimate trial-averaged activity and train the network, shown as a function of model dimensionality. (b) Cross-validated mean squared error (MSE) between the network’s firing rates and the trial-averaged activity computed on the held-out 20% of trials in each subset. Error bars indicate the standard error across folds (see Methods). Red traces correspond to networks trained on data from the left hemisphere, while blue traces correspond to networks trained on data from the right hemisphere.

**Figure S2:**
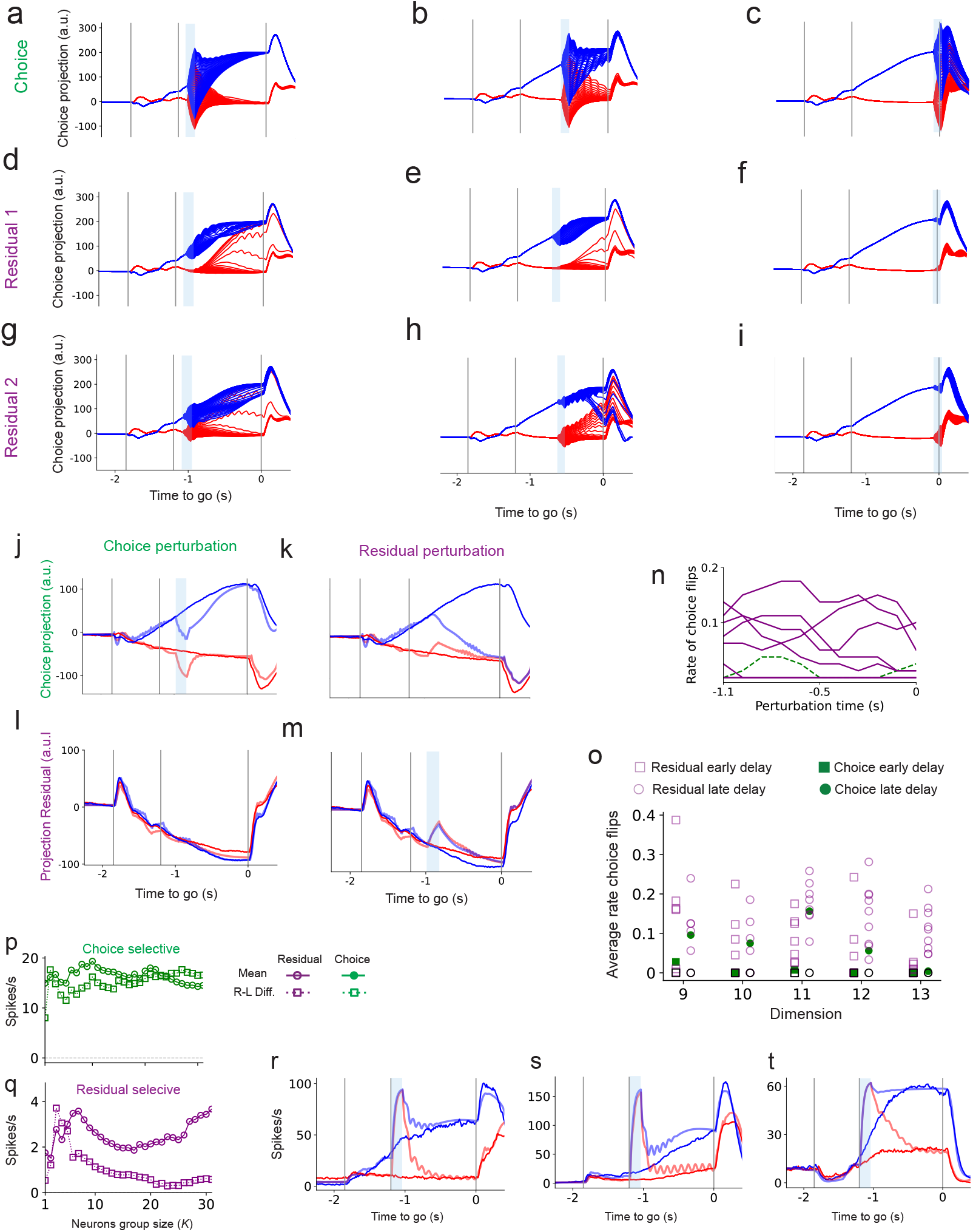
Supplementary Figure corresponding to Fig. 2. Projections onto the choice dimension for perturbations aligned with the choice dimension (a–c), the influential residual dimension from Fig. 2d,f (d–f), and the influential residual dimension from Fig. 2i (g–i), shown at the beginning, middle, and end of the delay epoch. All data in panels (a–i) correspond to the same network from the **Right Hemisphere** shown in Fig. 2 (*P* = 10). (j–m) Projections onto the choice (j,k) and residual (l,m) dimensions for perturbations aligned with the choice and residual directions, respectively, for a *different network trained with data from the* **Left Hemisphere**. The blue bar indicates the perturbation period. (n) Rate of choice flips for 11 distinct 100 ms perturbation intervals during the delay epoch across 10 network dimensions, using perturbation magnitudes from –15 to 15, for the Left Hemisphere network. (o) Extension of Fig. 2g showing average choice-flip rates from Fig. 2h, split into early (–1.1 to –0.6 s) and late (–0.5 to 0 s) delay epochs, summarized across networks trained with 9 to 13 dimensions. (p–q) Solid and circles: mean firing rate averaged over the delay epoch and over the top-*K* selective neurons, pooling left and right trials. Dashed and squares: mean right–minus–left firing-rate difference, averaged over the same delay window and neuron set. (p) choice-selective neurons; (q) residual-selective neurons. (r–t) Responses of the top three choice-selective neurons to perturbations targeting the top 30 choice-selective neurons using a positive perturbation (same magnitude and opposite sign to Fig. 2k).

**Figure S3:**
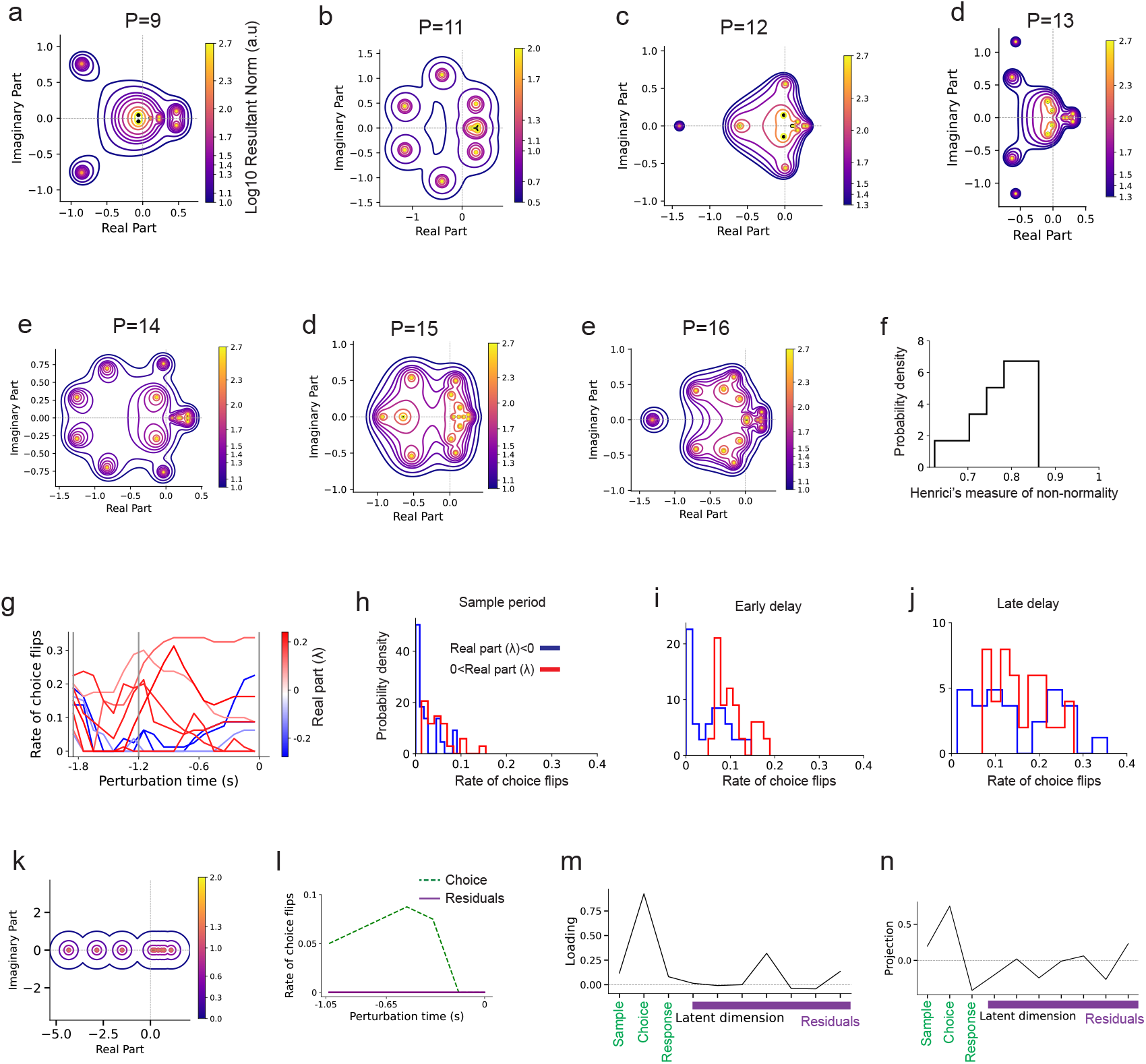
Supplementary Figure corresponding to Fig. 3. Eigenvalue spectra and pseudospectra of the interaction matrix (Trefethen & Embree 2020) for networks with dimensionality *P* = 9 (a) and *P* = 11–16 (b–e). (f) Normalized Henrici index (see Methods) across 15 trained networks with latent dimensionality ranging from P = 9 to P = 16 trained on neurons recorded on the left hemisphere. (g) Rate of choice flips for perturbations applied to each Schur dimension across 18 non-overlapping 100 ms windows spanning the sample and delay epochs. Each dimension was tested with perturbation magnitudes from –15 to 15. Line colors indicate the real part of the associated eigenvalue. (h–j) Mean rate of choice flips for Schur dimensions with negative (blue) and positive (red) real eigenvalues, averaged across perturbations applied during the sample epoch (h), early delay (–1.2 to –0.6 s; (i)), and late delay (–0.6 to 0 s; (j)). (k–n) Results for a network in which the interaction matrix was constrained to be symmetric (Methods), **A** = **A**^⊤^, with latent dimensionality *P* = 10. (k) Eigenvalue spectrum and pseudospectra of the interaction matrix, as in panels (a–e). The circular pseudospectral contours tightly centered on individual eigenvalues are the hallmark of normal (symmetric) connectivity, in contrast to the strongly balloon-like contours observed in the unconstrained networks (panels a–e), where the pseudospectrum extends far from the eigenvalues indicating transient amplification (Trefethen & Embree 2020). (l) Rate of choice flips for 8 distinct 150 ms perturbation intervals applied during the delay epoch, for perturbations along the choice dimension (dashed green) and the 7 residual dimensions (purple), as in Fig. 2g. In contrast to the unconstrained network, perturbations along the choice dimension are strongly effective while residual perturbations are largely ineffective, the reverse of what is observed experimentally (Daie et al. 2023). (m) Loadings of the eigenvector of **A** associated with the eigenvalue second closest to zero (λ ≈ 0.15; the closest eigenvalue, λ ≈ 0.11, does not show a dominant projection onto the choice dimension), projected onto the latent dimensions. The largest loading is onto the choice dimension, indicating that the symmetric network organizes near-integrator dynamics along the choice dimension. (n) Projection onto latent dimensions of the eigenvector of the Jacobian matrix, evaluated at the network trajectory during the first 100 ms of the delay epoch, associated with the eigenvalue closest to zero. The largest loading is onto the choice dimension, indicating that when the network dynamics are linearized around the early-delay trajectory, the choice dimension is the slowest mode, consistent with the symmetric network learning an approximate integrator along the choice dimension.

**Figure S4:**
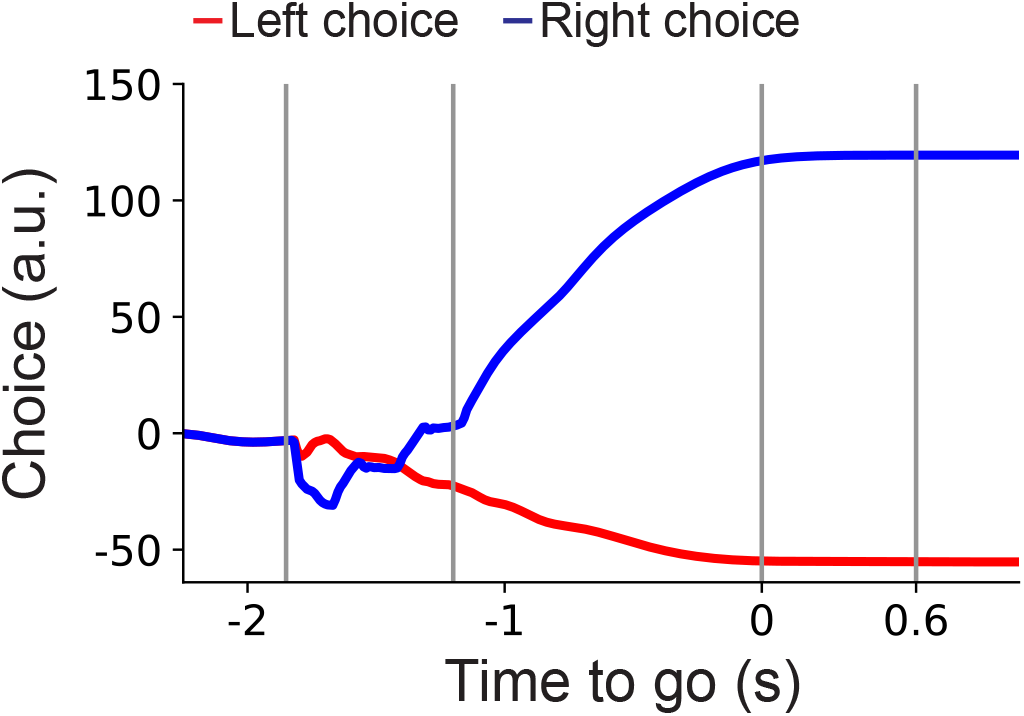
Supplementary Figure corresponding to Fig. 4. Projection onto the choice dimension for a network trained on ALM recordings from the left hemisphere (*P* = 13) when the go cue is withheld, illustrating convergence to a stable choice-selective attractors.

**Figure S5:**
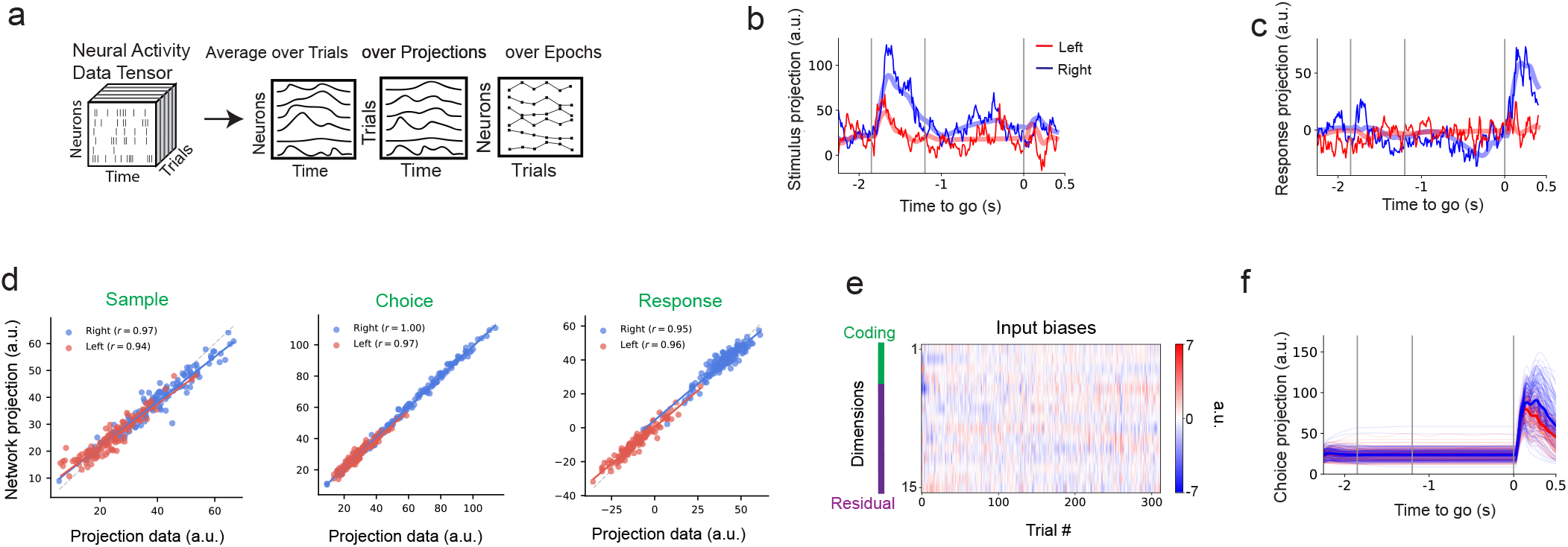
Supplementary Figure corresponding to Fig. 5. (a) Both the recurrent connectivity and trial-specific input biases (Eqs. (25, 26)) are optimized to fit the condition-averaged trajectories, their projections along coding dimensions, and the temporal average activity across all trial epochs (see Methods). (b–c) Single-trial dynamics projected onto the stimulus (b) and response (c) dimensions for left (red) and right (blue) trials. Solid lines represent recorded data; lighter lines show the corresponding network fits. (d) Comparison of neural trajectory projections onto the sample, choice, and response dimensions between the data and network model. Each point represents a single-trial, colored by choice direction (right, red; left, blue), with projections averaged over the sample epoch (sample dimension), delay epoch (choice dimension), and response epoch (response dimension), respectively. Regression lines shown per condition; Pearson correlation coefficients indicated in the legend. The diagonal dashed line denotes the identity. (e) Trial bias matrix after training, with *P* = 15 rows and columns corresponding to individual trials. (f) When the sample stimulus is withheld, the network fails to sustain delay-epoch activity, indicating that integration of the sample input is required to generate the preparatory activity. Input biases alone cannot account for the data, choice-encoding delay activity emerges only when the network integrates the sample stimulus.

**Figure S6:**
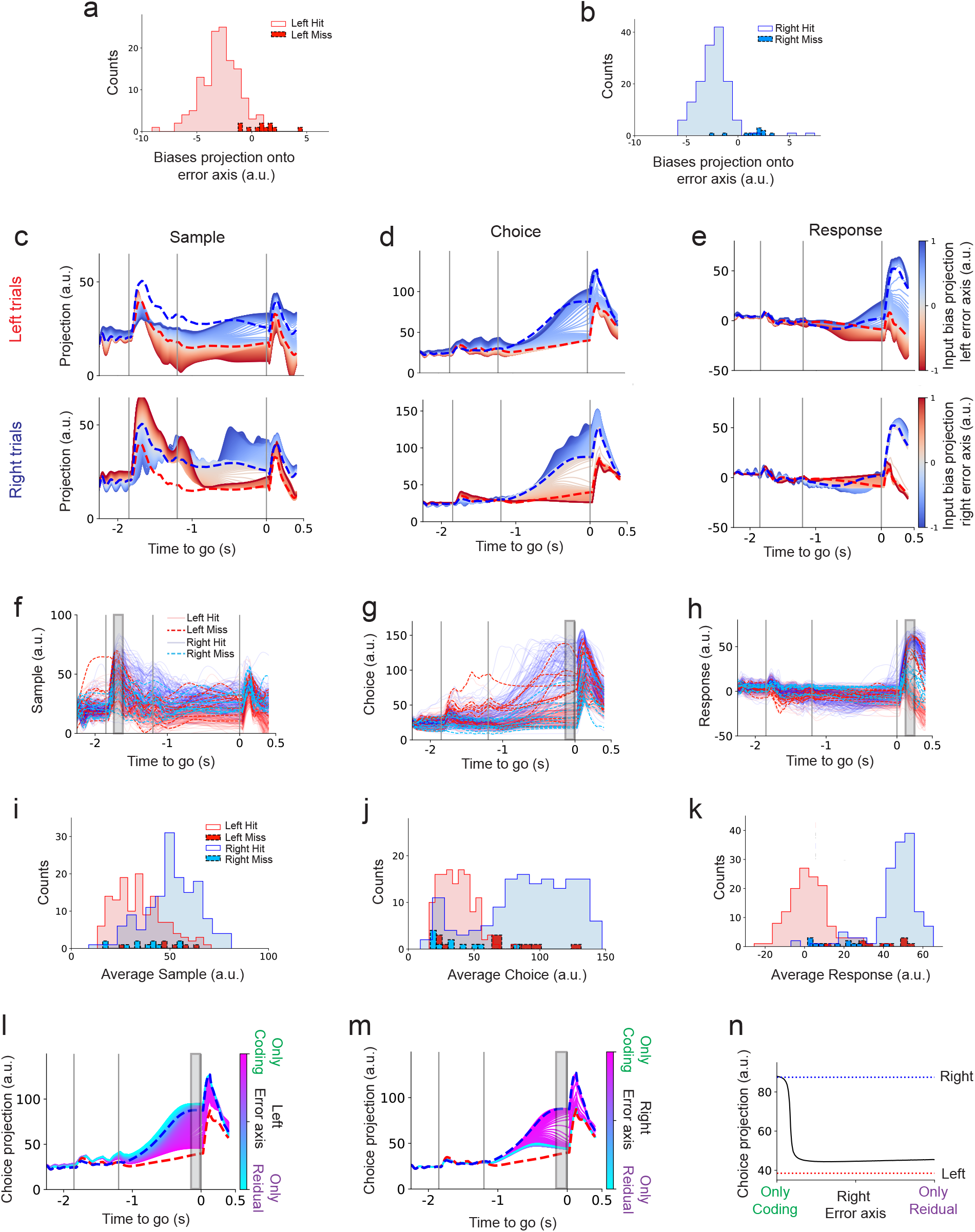
Supplementary Figure corresponding to Fig. 6. (a, b) Projections of input biases onto the left (a) and right (b) decoder weights (error axes) for correct and error trials in left (a) and right (b) conditions. (c–e) Network projections onto stimulus, choice, and response dimensions following perturbations along the error axes. Top: left stimulus trials; bottom: right stimulus trials. (f–h) Single-trial network trajectories projected onto stimulus, choice, and response dimensions for left (red) and right (blue) trials. Solid lines: correct trials; dashed lines: error trials. Deep red and cyan indicate left and right error trials, respectively. (i–k) Histograms of time-averaged projections (100 ms window; gray rectangles in f–h) across stimulus, choice, and response dimensions for each trial type. Trajectories in the choice projection for input biases of fixed magnitude that produce miss-like left (l) and right (m) trials (that is, trials in which the choice projection approaches the trial-averaged trajectory of right (l) and left (m) correct trials). The input bias magnitude is held constant while its composition is varied from input biases confined exclusively to coding dimensions (only coding) to those confined exclusively to residual dimensions (only residual) (see Methods). (n) Mean choice projection during the late delay epoch (215–235 ms; gray rectangle in panel (m)) plotted as a function of the relative contributions of coding and residual components in the input bias for the trials shown in (m).

**Figure S7:**
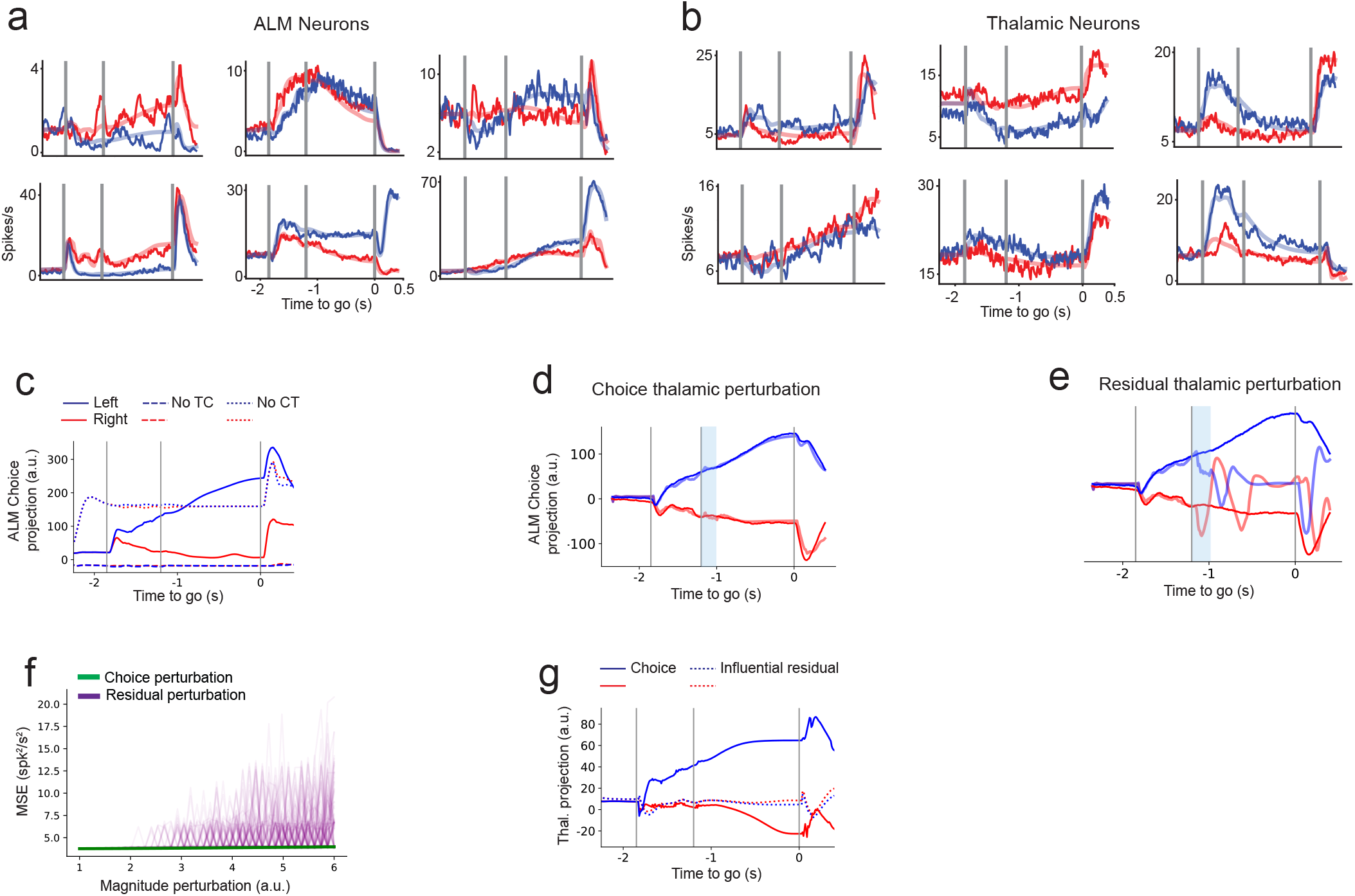
Supplementary Figure corresponding to Fig. 7. (a–b) Trial-averaged activity for correct left and right trials. Solid traces show neural recordings; lighter traces show model fits. Data are from the left hemisphere. Both cortex and thalamus were modeled with latent dimensionality *P* = 10, and both thalamocortical and corticothalamic projections were constrained to a bottleneck of 5 dimensions (see Methods). (a) ALM neurons; (b) VM and VL thalamic neurons. (c) Choice projection in ALM for three conditions: unperturbed (solid), thalamocortical projections removed (No TC, dashed), and corticothalamic projections removed (No CT, dotted). (d–e) Choice projection following perturbations in thalamus along the choice dimension (d) and a residual dimension (e). Yellow bar indicates the perturbation period. (f) Mean squared error (MSE) between unperturbed and perturbed trajectories for 120 perturbations along residual (purple) and choice (green) dimensions at increasing magnitudes. Light traces show individual perturbations; solid lines show the average. Panels (d–f) are from the left hemisphere and correspond to Fig. 7c–e. (g) Network thalamic activity projected onto the choice dimension (solid) and the influential residual dimension (dashed) used for the perturbation shown in Fig. 7d. The influential residual dimension exhibits little choice selectivity during the delay epoch compared with the choice dimension, yet perturbations along this dimension strongly disrupt cortical choice trajectories (Fig. 7d–e).

## References

Adesnik, H., & L. Abdeladim. 2021. “Probing neural codes with two-photon holographic optogenetics.” Nature Neuroscience 24: 1356–1366.

Ahrens, M. B., M. B. Orger, D. N. Robson, J. M. Li, & P. J. Keller. 2013. “Whole-brain functional imaging at cellular resolution using light-sheet microscopy.” Nature methods 10: 413–420.

Allen, W. E., M. Z. Chen, N. Pichamoorthy, R. H. Tien, M. Pachitariu, L. Luo, & K. Deisseroth. 2019. “Thirst regulates motivated behavior through modulation of brainwide neural population dynamics.” Science 364: eaav3932.

Amit, D. J., & N. Brunel. 1997. “Model of global spontaneous activity and local structured activity during delay periods in the cerebral cortex.” Cerebral cortex (New York, NY: 1991) 7: 237–252.

Angelaki, D., B. Benson, J. Benson, D. Birman, N. Bonacchi, K. Bougrova, S. A. Bruijns, M. Carandini, J. A. Catarino, et al. 2025. “A brain-wide map of neural activity during complex behaviour.” Nature 645: 177–191.

Asllani, M., R. Lambiotte, & T. Carletti. 2018. “Structure and dynamical behavior of non-normal networks.” Science Advances 4: eaau9403.

Barbosa, J., R. Proville, C. C. Rodgers, M. R. DeWeese, S. Ostojic, & Y. Boubenec. 2023. “Early selection of task-relevant features through population gating.” Nature Communications 14: 6837.

Ben-Yishai, R., R. L. Bar-Or, & H. Sompolinsky. 1995. “Theory of orientation tuning in visual cortex.” Proceedings of the National Academy of Sciences 92: 3844–3848.

Bounds, H. A., et al. 2024. “Network-level mechanisms of behaviorally silent populations.” bioRxiv.

Chen, G., B. Kang, J. Lindsey, S. Druckmann, & N. Li. 2021. “Modularity and robustness of frontal cortical networks.” Cell 184: 3717–3730.

Chen, S., Y. Liu, Z. A. Wang, J. Colonell, L. D. Liu, H. Hou, N.-W. Tien, et al. 2024. “Brain-wide neural activity underlying memory-guided movement.” Cell 187: 676–691.

Churchland, M. M., M. Y. Byron, M. Sahani, & K. V. Shenoy. 2007. “Techniques for extracting single-trial activity patterns from large-scale neural recordings.” Current opinion in neurobiology 17: 609–618.

Churchland, M. M., J. P. Cunningham, M. T. Kaufman, S. I. Ryu, & K. V. Shenoy. 2010. “Cortical preparatory activity: representation of movement or first cog in a dynamical machine?” Neuron 68: 387–400.

Clark, D. G., & M. Beiran. 2024. “Structure of activity in multiregion recurrent neural networks.” arXiv preprint 2402.12188.

Compte, A., N. Brunel, P. S. Goldman-Rakic, & X.-J. Wang. 2000. “Synaptic mechanisms and network dynamics underlying spatial working memory in a cortical network model.” Cerebral cortex 10: 910–923.

Cunningham, J. P., & B. M. Yu. 2014. “Dimensionality reduction for large-scale neural recordings.” Nature Neuroscience 17: 1500–1509.

Daie, K., L. Fontolan, S. Druckmann, & K. Svoboda. 2023. “Feedforward amplification in recurrent networks underlies paradoxical neural coding.” bioRxiv, 2023–08.

Daie, K., K. Svoboda, & S. Druckmann. 2021. “Targeted photostimulation uncovers circuit motifs supporting shortterm memory.” Nature Neuroscience 24: 259–265.

Darshan, R., & A. Rivkind. 2022. “Learning to represent continuous variables in heterogeneous neural networks.” Cell Reports 39.

DePasquale, B., D. Sussillo, L. Abbott, & M. M. Churchland. 2023. “The centrality of population-level factors to network computation is demonstrated by a versatile approach for training spiking networks.” Neuron 111: 631–649.

Elsayed, G. F., A. H. Lara, M. T. Kaufman, M. M. Churchland, & J. P. Cunningham. 2016. “Reorganization between preparatory and movement population responses in motor cortex.” Nature Communications 7: 13239.

Finkelstein, A., L. Fontolan, M. N. Economo, N. Li, S. Romani, & K. Svoboda. 2021. “Attractor dynamics gate cortical information flow during decision-making.” Nature Neuroscience 24: 843–850.

Gallego, J. A., M. G. Perich, L. E. Miller, & S. A. Solla. 2017. “Neural manifolds for the control of movement.” Neuron 94: 978–984.

Ganguli, S., D. Huh, & H. Sompolinsky. 2008. “Memory traces in dynamical systems.” Proceedings of the national academy of sciences 105: 18970–18975.

Gao, Z., C. Davis, A. M. Thomas, M. N. Economo, A. M. Abrego, K. Svoboda, C. I. De Zeeuw, & N. Li. 2018. “A cortico-cerebellar loop for motor planning.” Nature 563: 113–116.

Gauld, C., et al. 2024. “Latent, task-silent cortical states shape behavior through inhibition and recruitment.” Nature.

Georgopoulos, A. P., R. E. Kettner, & A. B. Schwartz. 1988. “Primate motor cortex and free arm movements to visual targets in three-dimensional space. II. Coding of the direction of movement by a neuronal population.” Journal of Neuroscience 8: 2928–2937.

Gillett, M., & N. Brunel. 2024. “Dynamic control of sequential retrieval speed in networks with heterogeneous learning rules.” eLife 12: RP88805.

Gillett, M., U. Pereira, & N. Brunel. 2020. “Characteristics of sequential activity in networks with temporally asymmetric Hebbian learning.” Proceedings of the National Academy of Sciences 117: 29948–29958.

Gold, J. I., & M. N. Shadlen. 2007. “The neural basis of decision making.” Annu. Rev. Neurosci. 30: 535–574.

Goldman, M. S. 2009. “Memory without feedback in a neural network.” Neuron 61: 621–634.

Golub, G. H., & C. F. Van Loan. 2013. Matrix computations. JHU press.

Grossberg, S. 1969. “On learning, information, lateral inhibition, and transmitters.” Mathematical Biosciences 4: 255– 310.

Guckenheimer, J., & P. Holmes. 2013. Nonlinear oscillations, dynamical systems, and bifurcations of vector fields. Vol. 42. Springer Science & Business Media.

Guo, Z. V., et al. 2017. “Maintenance of persistent activity in a frontal thalamocortical loop.” Nature 545: 181–186.

Guo, Z. V., N. Li, D. Huber, E. Ophir, D. Gutnisky, J. T. Ting, G. Feng, & K. Svoboda. 2014. “Flow of Cortical Activity Underlying a Tactile Decision in Mice.” Neuron 81: 179–194.

Hennequin, G., T. P. Vogels, & W. Gerstner. 2012. “Nonnormal amplification in random balanced neuronal networks.” Physical Review E—Statistical, Nonlinear, and Soft Matter Physics 86: 011909.

Hopfield, J. J. 1982. “Neural networks and physical systems with emergent collective computational abilities.” Proceedings of the National Academy of Sciences 79: 2554–2558.

Hopfield, J. J. 1984. “Neurons with graded response have collective computational properties like those of two-state neurons.” Proceedings of the national academy of sciences 81: 3088–3092.

Inagaki, H. K., S. Chen, M. C. Ridder, P. Sah, N. Li, Z. Yang, H. Hasanbegovic, et al. 2022. “A midbrain-thalamus-cortex circuit reorganizes cortical dynamics to initiate movement.” Cell 185: 1065–1081.

Inagaki, H. K., L. Fontolan, S. Romani, & K. Svoboda. 2019. “Discrete attractor dynamics underlies persistent activity in the frontal cortex.” Nature 566: 212–217.

Inagaki, H. K., M. Inagaki, S. Romani, & K. Svoboda. 2018. “Low-dimensional and monotonic preparatory activity in mouse anterior lateral motor cortex.” Journal of Neuroscience 38: 4163–4185.

Jun, J. J., N. A. Steinmetz, J. H. Siegle, D. J. Denman, M. Bauza, B. Barbarits, A. K. Lee, et al. 2017. “Fully integrated silicon probes for high-density recording of neural activity.” Nature 551: 232–236.

Kaku, H., L. D. Liu, R. Gao, S. West, S.-M. Liao, A. Finkelstein, D. Kleinfeld, et al. 2025. “A brainstem map of orofacial rhythms.” bioRxiv.

Kaufman, M. T., et al. 2014. “Cortical activity in the null space: permitting preparation without movement.” Nature Neuroscience 17: 440–448.

Khilkevich, A., et al. 2024. “Beyond preparatory activity: motor cortex dynamics in subspaces not aligned with movement.” bioRxiv.

Khona, M., & I. R. Fiete. 2022. “Attractor and integrator networks in the brain.” Nature Reviews Neuroscience 23: 744– 766.

Kim, C. M., A. Finkelstein, C. C. Chow, K. Svoboda, & R. Darshan. 2023. “Distributing task-related neural activity across a cortical network through task-independent connections.” Nature Communications 14: 2851.

Kleinfeld, D. 1986. “Sequential state generation by model neural networks.” Proceedings of the National Academy of Sciences 83: 9469–9473.

Kobak, D., W. Brendel, C. Constantinidis, C. E. Feierstein, A. Kepecs, Z. F. Mainen, X.-L. Qi, et al. 2016. “Demixed principal component analysis of neural population data.” eLife 5: e10989.

Langdon, C., & T. A. Engel. 2025. “Latent circuit inference from heterogeneous neural responses during cognitive tasks.” Nature Neuroscience, 1–11.

Li, N., K. Daie, K. Svoboda, & S. Druckmann. 2016. “Robust neuronal dynamics in premotor cortex during motor planning.” Nature 532: 459–464.

Luo, T. Z., T. D. Kim, D. Gupta, A. G. Bondy, C. D. Kopec, V. A. Elliott, B. DePasquale, & C. D. Brody. 2025. “Transitions in dynamical regime and neural mode during perceptual decisions.” Nature, 1–11.

MacDowell, C. J., A. Libby, C. I. Jahn, S. Tafazoli, A. Ardalan, & T. J. Buschman. 2025. “Multiplexed subspaces route neural activity across brain-wide networks.” Nature Communications 16: 3359.

Maimon, G., & J. A. Assad. 2006. “A cognitive signal for the proactive timing of action in macaque LIP.” Nature neuroscience 9: 948–955.

Mante, V., D. Sussillo, K. V. Shenoy, & W. T. Newsome. 2013. “Context-dependent computation by recurrent dynamics in prefrontal cortex.” Nature 503: 78–84.

Marshel, J. H., Y. S. Kim, T. A. Machado, S. Quirin, B. Benson, J. Kadmon, C. Raja, et al. 2019. “Cortical layer– specific critical dynamics triggering perception.” Science 365: eaaw5202.

Mastrogiuseppe, F., & S. Ostojic. 2018. “Linking connectivity, dynamics, and computations in low-rank recurrent neural networks.” Neuron 99: 609–623.

Matteucci, G., M. Guyoton, J. M. Mayrhofer, M. Auffret, G. Foustoukos, C. C. Petersen, & S. El-Boustani. 2022. “Cortical sensory processing across motivational states during goal-directed behavior.” Neuron 110: 4176–4193.

Miller, K. D., & F. Fumarola. 2012. “Mathematical equivalence of two common forms of firing rate models of neural networks.” Neural Computation 24: 25–31.

Monsalve-Mercado, M. M., & K. D. Miller. 2025. “The geometry of the neural state space of decisions.” bioRxiv, 2025– 01.

Murphy, B. K., & K. D. Miller. 2009. “Balanced amplification: a new mechanism of selective amplification of neural activity patterns.” Neuron 61: 635–648.

O’Shea, D. J., L. Duncker, W. Goo, X. Sun, S. Vyas, E. M. Trautmann, I. Diester, et al. 2022. “Direct neural perturbations reveal a dynamical mechanism for robust computation.” bioRxiv, 2022–12.

Oña-Jodar, T., G. Prat-Ortega, C. Li, J. Dalmau, A. Compte, & J. de la Rocha. 2024. “Episodic recruitment of attractor dynamics in frontal cortex reveals distinct mechanisms for forgetting and lack of cognitive control in short-term memory.” bioRxiv, 2024–02.

Orhan, A. E., & W. J. Ma. 2019. “A diverse range of factors affect the nature of neural representations underlying short-term memory.” Nature neuroscience 22: 275–283.

Pals, M., A. E. Sagtekin, F. Pei, M. Gloeckler, & J. H. Macke. 2024. Inferring stochastic low-rank recurrent neural networks from neural data.

Pang, R., B. J. Lansdell, & A. L. Fairhall. 2016. “Dimensionality reduction in neuroscience.” Current Biology 26: R656– R660.

Pellegrino, A., H. Stein, & N. A. Cayco-Gajic. 2024. “Dimensionality reduction beyond neural subspaces with slice tensor component analysis.” Nature Neuroscience 27: 1199– 1210.

Pereira-Obilinovic, U., S. Froudist-Walsh, & X.-J. Wang. 2024. “Cognitive network interactions through communication subspaces in large-scale models of the neocortex.” bioRxiv.

Perich, M. G., D. Narain, & J. A. Gallego. 2025. “A neural manifold view of the brain.” Nature Neuroscience 28: 1582– 1597.

Poulet, J. F., & C. C. Petersen. 2008. “Internal brain state regulates membrane potential synchrony in barrel cortex of behaving mice.” Nature 454: 881–885.

Qian, W., J. Zavatone-Veth, B. Ruben, & C. Pehlevan. 2024. “Partial observation can induce mechanistic mismatches in data-constrained models of neural dynamics.” Advances in Neural Information Processing Systems 37: 67467–67510.

Rajan, K., C. D. Harvey, & D. W. Tank. 2016. “Recurrent network models of sequence generation and memory.” Neuron 90: 128–142.

Recanatesi, S., U. Pereira-Obilinovic, M. Murakami, Z. Mainen, & L. Mazzucato. 2022. “Metastable attractors explain the variable timing of stable behavioral action sequences.” Neuron 110: 139–153.

Robinson, N. T., L. A. Descamps, L. E. Russell, M. O. Buchholz, B. A. Bicknell, G. K. Antonov, J. Y. Lau, et al. 2020. “Targeted activation of hippocampal place cells drives memory-guided spatial behavior.” Cell 183: 1586–1599.

Salzman, C. D., K. H. Britten, & W. T. Newsome. 1990. “Cortical microstimulation influences perceptual judgements of motion direction.” Nature 346: 174–177.

Semedo, J. D., et al. 2019. “Cortical areas interact through a communication subspace.” Neuron 102: 249–259.

Seung, H. S. 1996. “How the brain keeps the eyes still.” Proceedings of the National Academy of Sciences 93: 13339–13344.

Sofroniew, N. J., D. Flickinger, J. King, & K. Svoboda. 2016. “A large field of view two-photon mesoscope with subcellular resolution for in vivo imaging.” elife 5: e14472.

Sompolinsky, H., & I. Kanter. 1986. “Temporal association in asymmetric neural networks.” Physical Review Letters 57: 2861.

Sourmpis, C., C. Petersen, W. Gerstner, & G. Bellec. 2023. “Trial matching: capturing variability with data-constrained spiking neural networks.” Advances in Neural Information Processing Systems 36: 74787–74798.

Sourmpis, C., C. C. Petersen, W. Gerstner, & G. Bellec. 2024. “Biologically informed cortical models predict optogenetic perturbations.” bioRxiv, 2024–09.

Steinemann, N., G. M. Stine, E. Trautmann, A. Zylberberg, D. M. Wolpert, & M. N. Shadlen. 2024. “Direct observation of the neural computations underlying a single decision.” eLife 12: RP90859.

Steinmetz, N. A., P. Zatka-Haas, M. Carandini, & K. D. Harris. 2019. “Distributed coding of choice, action and engagement across the mouse brain.” Nature 576: 266–273.

Stopfer, M., V. Jayaraman, & G. Laurent. 2003. “Intensity versus identity coding in an olfactory system.” Neuron 39: 991–1004.

Stroud, J. P., J. Duncan, & M. Lengyel. 2024. “The computational foundations of dynamic coding in working memory.” Trends in cognitive sciences 28: 614–627.

Stroud, J. P., K. Watanabe, T. Suzuki, M. G. Stokes, & M. Lengyel. 2023. “Optimal information loading into working memory explains dynamic coding in the prefrontal cortex.” Proceedings of the National Academy of Sciences 120: e2307991120.

Tanaka, M. 2007. “Cognitive signals in the primate motor thalamus predict saccade timing.” Journal of Neuroscience 27: 12109–12118.

Tanji, J., & E. V. Evarts. 1976. “Anticipatory activity of motor cortex neurons in relation to direction of an intended movement.” Journal of Neurophysiology 39: 1062–1068.

Thura, D., & P. Cisek. 2014. “Deliberation and commitment in the premotor and primary motor cortex during dynamic decision making.” Neuron 81: 1401–1416.

Trefethen, L. N., & M. Embree. 2020. “Spectra and pseudospectra: the behavior of nonnormal matrices and operators.”

Valente, A., J. W. Pillow, & S. Ostojic. 2022. “Extracting computational mechanisms from neural data using low-rank RNNs.” Advances in Neural Information Processing Systems 35: 24072–24086.

Vyas, S., et al. 2020. “Computation through neural population dynamics.” Annual Review of Neuroscience 43: 249–275.

Wang, X.-J. 2002. “Probabilistic decision making by slow reverberation in cortical circuits.” Neuron 36: 955–968.

Wang, Y., X. Yin, Z. Zhang, J. Li, W. Zhao, & Z. V. Guo. 2021. “A cortico-basal ganglia-thalamo-cortical channel underlying short-term memory.” Neuron 109: 3486–3499.

Williams, A. H., T. H. Kim, F. Wang, S. Vyas, S. I. Ryu, K. V. Shenoy, M. Schnitzer, T. G. Kolda, & S. Ganguli. 2018. “Unsupervised discovery of demixed, low-dimensional neural dynamics across multiple timescales through tensor component analysis.” Neuron 98: 1099–1115.

Wimmer, K., A. Compte, A. Roxin, D. Peixoto, A. Renart, & J. De La Rocha. 2015. “Sensory integration dynamics in a hierarchical network explains choice probabilities in cortical area MT.” Nature Communications 6: 6177.

Wong, K.-F., & X.-J. Wang. 2006. “A recurrent network mechanism of time integration in perceptual decisions.” Journal of Neuroscience 26: 1314–1328.

Yang, H., et al. 2022. “Distinct thalamocortical pathways for maintaining and updating working memory.” Nature 609: 113–121.

Yang, W., S. L. Tipparaju, G. Chen, & N. Li. 2022. “Thalamus-driven functional populations in frontal cortex support decision-making.” Nature neuroscience 25: 1339–1352.

